# Deforming the metric of cognitive maps distorts memory

**DOI:** 10.1101/391201

**Authors:** Jacob L. S. Bellmund, William de Cothi, Tom A. Ruiter, Matthias Nau, Caswell Barry, Christian F. Doeller

## Abstract

Environmental boundaries anchor cognitive maps that support memory. However, trapezoidal boundary geometry distorts the regular firing patterns of entorhinal grid cells proposedly providing a metric for cognitive maps. Here, we test the impact of trapezoidal boundary geometry on human spatial memory using immersive virtual reality. Consistent with reduced regularity of grid patterns in rodents and a grid-cell model based on the eigenvectors of the successor representation, human positional memory was degraded in a trapezoid compared to a square environment; an effect particularly pronounced in the trapezoid’s narrow part. Congruent with spatial frequency changes of eigenvector grid patterns, distance estimates between remembered positions were persistently biased; revealing distorted memory maps that explained behavior better than the objective maps. Our findings demonstrate that environmental geometry affects human spatial memory similarly to rodent grid cell activity — thus strengthening the putative link between grid cells and behavior along with their cognitive functions beyond navigation.

## Introduction

Boundaries are essential for navigators moving through space. Consistently, boundary geometry serves as a strong cue for reorientation (Cheng and Newcombe, 2005; Julian et al., 2018). When rodents and human children are disoriented after learning the location of a hidden reward, they search for the reward equally often in geometrically equivalent corners of rectangular environments (Cheng, 1986; Margules and Gallistel, 1988; Hermer and Spelke, 1994). Consistently, human adults rely on boundary geometry for spatial updating, facilitated by a limited number of symmetry axes of enclosure boundaries (Kelly et al., 2008). The learning of positions relative to a boundary, which recruits the hippocampal formation, is thought to occur incidentally (Doeller and Burgess, 2008; Doeller et al., 2008) and positions closer to a boundary are remembered more accurately than those further away from it (Lee et al., 2018).

Here, we examine the possibility that boundary geometry can cause distortions in human spatial memory. We derive this hypothesis from the distortions induced by environmental geometry on rodent grid-cell firing patterns (Krupic et al., 2015, 2018; Stensola et al., 2015). Grid cells, first identified in the entorhinal cortex of freely moving rodents, typically exhibit six-fold periodic (hexadirectional) spatial firing extending across the environment (Hafting et al., 2005). This pattern can be described in terms of its scale, as well as its offset and orientation relative to the environment (Hafting et al., 2005; Moser et al., 2017). Along the dorso-ventral axis of medial entorhinal cortex, grid cells sharing similar spacing and orientations are organized in discrete modules (Barry et al., 2007; Brun et al., 2008; Stensola et al., 2012). Grid cells have been directly recorded in human patients undergoing pre-surgical screening (Jacobs et al., 2013; Nadasdy et al., 2017) and in human fMRI studies hexadirectional signals serve as a proxy measure for activity of the entorhinal grid system (Doeller et al., 2010). However, empirical evidence demonstrating the behavioral relevance of grid cells remains scarce.

Theoretical work suggests that regular grid patterns provide a compact code for self-localization and function as a metric for space, supporting path integration and vector-based navigation (Moser et al., 2017; Hafting et al., 2005; McNaughton et al., 2006; Fiete et al., 2008; Burak and Fiete, 2009; Mathis et al., 2012; Bush et al., 2015; Herz et al., 2017; Banino et al., 2018). Thus, location is encoded by the conjunction of spatial phases across different modules — the population phase (Bush et al., 2015; Carpenter and Barry, 2016) — while the distance and direction between points can be derived from the relative difference in population phase (Bush et al., 2015). With a regular underlying grid pattern, there should be a tight coupling between the distance separating positions and the change in grid population phase. Larger distances in space correspond to greater changes in grid population phase.

Environmental geometry strongly influences grid firing patterns in rodents (Krupic et al., 2015; Stensola et al., 2015; Sun et al., 2015; Krupic et al., 2018). Changes made to the geometry of a familiar enclosure produce commensurate changes to the scale of grid-patterns, resulting in differential rates of change in population phase for travel in the changed and unchanged dimension (Barry et al., 2007; Stensola et al., 2012). Similar manipulations made while humans navigate in VR environments produce complementary deficits in path integration (Chen et al., 2015). More strikingly, in highly polarized enclosures such as trapezoids, grid-patterns are highly distorted and less regular than in control enclosures (Krupic et al., 2015). These changes are especially pronounced in the narrow part of the trapezoidal enclosure with reduced symmetry, less regular fields and a change of grid orientation — changes which do not appear to attenuate with continued exposure (Krupic et al., 2015). Similarly, in a quadrilateral environment with one slanted wall, firing fields of grid cells were consistently shifted away from the slanted wall, resulting in a local distortion of the grid (Krupic et al., 2018). Together, these findings indicate that environmental boundaries not only anchor spatial representations, but that their specific arrangement can distort spatial codes in the mammalian brain. However, the potential consequences of compromised grid patterns for human spatial cognition have not been explored.

Here, we investigate how environmental geometry — which is known to distort grid-cell based computations — influences human spatial cognition. Degraded grid patterns in a trapezoid are hypothesized to carry less precise positional information than regular grid patterns, resulting in uncertainties about locations in space and the distances between them (Carpenter and Barry, 2016; Krupic et al., 2015, 2018). Thus, we investigated the effects of boundary geometry on human spatial memory. We capture the effects of environmental geometry on grid patterns using the eigenvectors of the successor representation (SR, see Methods), which exhibit grid-like properties in two-dimensional space whose regularity is degraded in a trapezoid (Stachenfeld et al., 2017). We demonstrate that distorted eigenvector grid patterns convey less precise information about self-location in a trapezoid, this effect being most pronounced in the narrow end. We tested whether memory for object positions is impaired in a trapezoid compared to a square control environment. Within the trapezoid, we expected worse memory performance particularly in the narrow compared to the broad part of the enclosure. In addition, spatial computations performed on the basis of distorted grid patterns are expected to exhibit systematic biases. SR grid patterns were, on average, stretched in the trapezoid relative to the square and compressed in the narrow compared to broad part of the trapezoid. We asked participants to judge distances between remembered locations and contrasted their estimates of identical true distances as a function of environmental geometry.

## Results

### Positional memory

We employed immersive VR to investigate effects of environmental geometry on human spatial memory (Figure 1A). Wearing a head-mounted display, participants navigated different environments using a motion platform translating real-world rotations and steps into virtual movement (Figure 1B). Participants were familiarized with the VR setup in a circular environment before learning object positions in a square and a trapezoid with the order of environments counterbalanced across participants. The environments were of equal surface area and distinct wall colors served as orientation cues. In the object position memory task, participants learned the positions of six objects in each environment, organized in two triplets with matched inter-object distances in both halves of an environment (Figure 1C). Participants were tested on the positions of the objects after an initial learning phase by having to navigate to the remembered position of a cued object in each trial (Figure 1D). To probe mnemonic distortions outside of the encoding environment, participants judged pairwise distances between object positions in VR by walking the distance in the circular familiarization environment and on a computer screen by adjusting a slider on a subjective scale.

**Figure 1.**
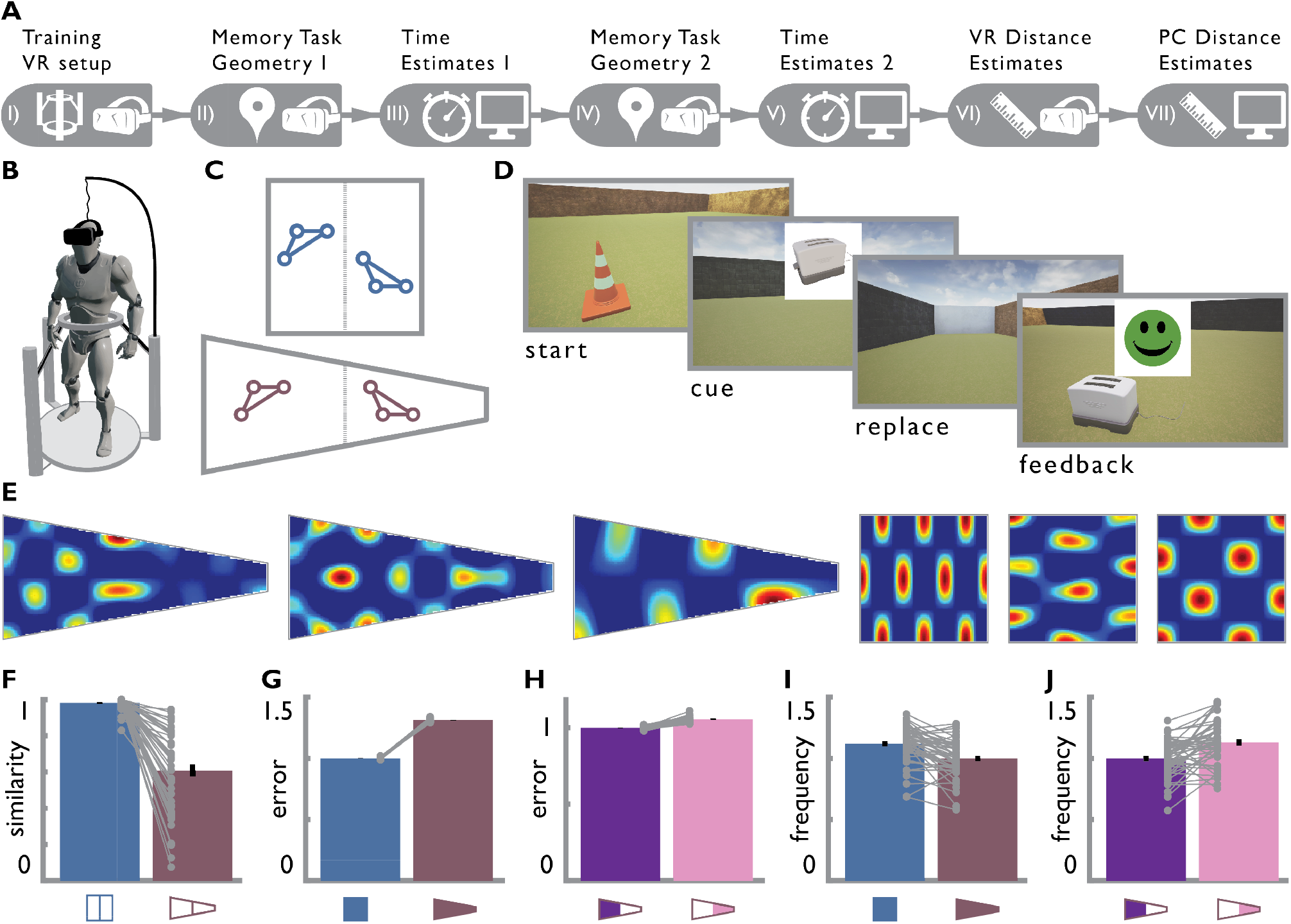
Task design and SR grid patterns. **A.** Experimental timeline. Participants were (I) familiarized with the VR setup before completing the (II & IV) object position memory task followed by (III & V) time estimates in the two environments. Afterwards (VI & VII) participants estimated pairwise distances between learned positions in VR and on a computer screen. Headset and screen icon indicate whether individual tasks took place in VR or on a computer screen, respectively. **B.** Schematic of immersive VR setup with head mounted display and motion platform translating physical steps and rotations into virtual movement. **C.** Circles illustrate an exemplary configuration of object positions. Two triplets of objects were positioned in each environment with one triplet in each half of each environment, yielding four triplets with matched distances between positions. **D.** To commence a test trial in the object position memory task, participants walked to a start position marked by a pylon where they were cued with the image of an object. Subsequently, they navigated to the object’s remembered position, which they indicated via button press, and received feedback. **E.** Example eigenvector grid patterns of the successor representation for a trapezoid (top) and a square (bottom) environment (Supplemental Figure 1). **F.** SR grid patterns are distorted in the trapezoid resulting in reduced correlation coefficients of spatial autocorrelations between the two trapezoid halves compared to the halves of the square. **G, H.** Position decoding errors based on spikes sampled from SR grid patterns are (**G**) larger in the trapezoid than in the square and (**H**) larger in the narrow than the broad part of the trapezoid, demonstrating that distorted grid patterns carry less positional information (Supplemental Figure 2). **I, J.** Mean radial frequencies of SR grid patterns are (**I**) lower in the trapezoid than in the square and (**J**) higher in the narrow than the broad part of the trapezoid (Supplemental Figure 3). Bars in **F-J** show mean and SEM. Individual data points reflect iterations (**G, H**) or SR grid patterns (**F, I, J**).

Does the disruption of regular grid patterns (Figure 1EF, Supplemental Figure 1, t(98)=10.81, p<0.001, d= 2.11, confidence interval CI: 0.31, 0.45) result in less accurate positional codes? To test this notion, we used a Bayesian decoder to decode locations using synthetic spike trains sampled from a population of SR grid patterns (n=50), themselves derived from the eigenvectors of successor representations from the square and trapezoid environment. Decoding was performed using a simple maximum likelihood approach assuming uniform priors (Towse et al., 2014). Decoding errors — the displacement between the true and decoded position — were larger in the trapezoid than in the square (Figure 1G, t(58)=117.41, p<0.001,Cohen’s d=31.36, CI: 0.31; 0.32), indicating that distorted grid patterns carry less positional information. To exclude the possibility that this reduction simply reflected the change in environmental aspect ratio, we verified that decoding accuracy for the smallest rectangular environment entirely enclosing the trapezoid exceeds that for the trapezoid itself (Supplemental Figure 2, t(58)=64.52, p<0.001, d=16.62, CI: 0.15; 0.16). Thus, distorted grid patterns indeed underlie worse position decoding in the trapezoid.

Is human positional memory also degraded in a trapezoid? In a first step, we compared raw positional memory error — the displacement between the response and correct position — between the two environments. Consistent with the degradation in positional information seen in simulated grid patterns, participants made larger errors in the trapezoid than the square (Figure 2A, t(36)=2.71, p<0.001, d=0.45, CI: 0.41; 2.06; bootstrap-based t-tests and confidence intervals are reported throughout the manuscript, see Methods). To ensure that this effect was not due to the fact that the trapezoid allows larger errors because of its elongated shape, we calculated memory scores accounting for differences in the distribution of possible errors for each position (Jacobs et al., 2016). We generated a chance distribution of 1000 random locations uniformly covering the entire environment and quantified the distance of each random location to the correct positions, resulting in a specific distribution of possible error distances for each position. For each trial, we calculated the memory score as 1 minus the proportion of distances from the chance distribution smaller than the replacement error. This yielded a score ranging from 0 (low memory) to 1 (perfect memory) for each trial, taking into account the range of possible errors based on the correct position and environmental geometry (see Supplemental Figure 4A for the overall distribution of memory scores). Importantly, memory scores were significantly lower in the trapezoid compared to the square (Figure 2BC, t(36)=-2.30, p<0.001, d=-0.38, CI: −0.04; −0.01), ensuring that decreased positional memory was not due to different distributions of possible errors as a result of the elongated shape of the trapezoid.

**Figure 2.**
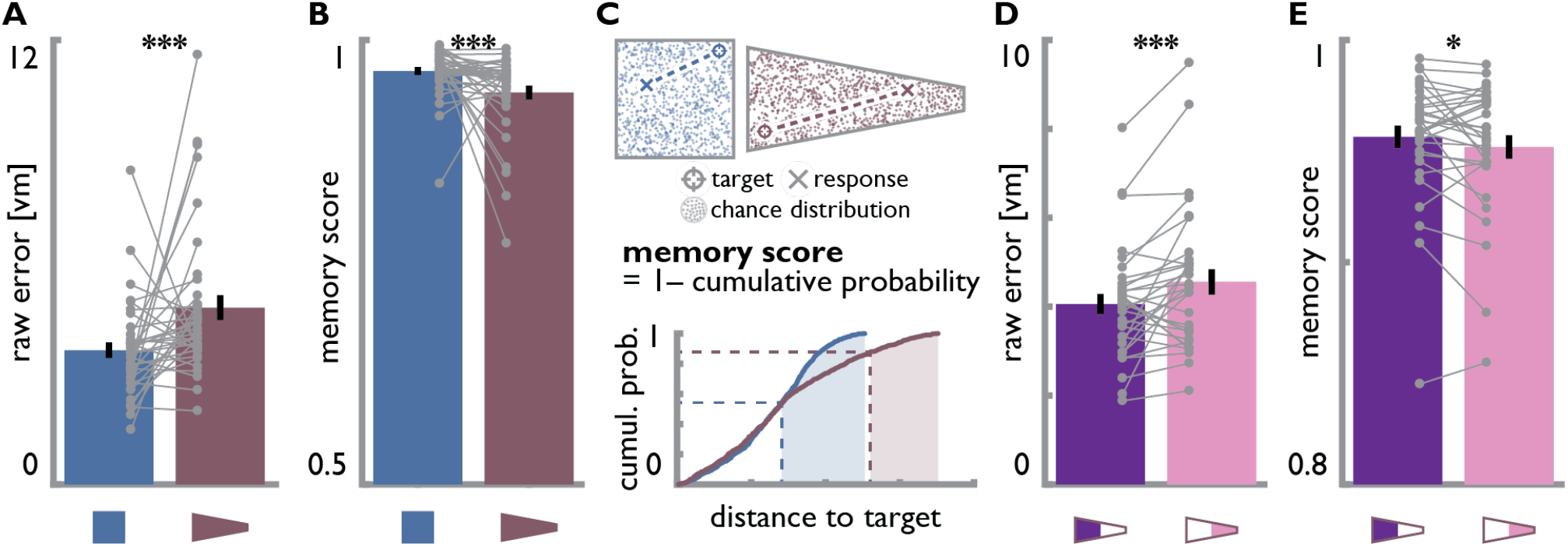
Distorted positional memory in the trapezoid. **A.** Positional memory errors in virtual meters (vm), measured as the Euclidean distance between the correct and remembered position, were larger for object positions in the trapezoid than in the square. **B.** Similarly, memory scores were lower in the trapezoid than in the square environment. Y-axis thresholded at chance level of 0.5. **C.** Schematics illustrate the expression of positional memory using memory scores to control for differences in distributions of possible errors for two hypothetical trials. For both environments, a chance distribution of positions uniformly covering the available space was generated (top, colored dots). We quantified the cumulative probability of the chance distribution as a function of distance to target object positions (bottom). Positional memory performance of a given trial was then expressed as 1-the cumulative probability at the error distance between the true and response position, resulting in memory scores ranging from 0-1, where 0.5 constitutes chance level and 1 perfect performance. Conceptually, this corresponds to the proportion of the chance distribution further away from the correct position than the remembered position (shaded areas). **D.** Raw positional memory errors were larger in the narrow compared to the broad part of the trapezoid. 5 participants were excluded due to differences between broad and narrow part more than 1.5 times the interquartile range above or below the upper and lower quartile of differences respectively. See Supplemental Figure 4B for full dataset. **E.** Likewise, memory scores were lower in the narrow than the broad part of the trapezoid. Y-axis thresholded to illustrate the difference between conditions. All bars show mean±SEM, grey circles indicate individual subject data with lines connecting data points from the same participant. * p<0.05, *** p<0.001

In rodents, grid patterns recorded from a trapezoid are known to be more strongly distorted in the narrow end of the environment than its base (Krupic et al., 2015). Similarly, we found that decoding errors derived from SR grid patterns were also larger in the narrow part of the trapezoid (Figure 1H, t(58)=14.63, p<0.001, d=3.83, CI: 0.05; 0.07). To determine whether human spatial memory differed within the trapezoid we examined memory errors finding, as expected, that errors were larger in the narrow end (Figure 2D, t(31)=2.75, p<0.001, d=0.49, CI: 0.18; 0.87). To control for the expected difference in error distributions we again calculated memory scores, confirming a robust difference between the two ends of the environment (Figure 2E, t(31)=-1.59, p=0.023, d=-0.28, CI: −0.01; 0.0006). The difference in positional memory errors between the narrow and broad part of the trapezoid was higher than the 5^th^ percentile of a surrogate distribution of error differences obtained from comparing positional memory between the halves of the square, indicating a significant difference between the environments (Supplemental Figure 4C, Z=2.18, p=0.015). Taken together, the profile of positional memory observed is in line with our predictions derived from deformations of grid patterns with degraded positional memory in the trapezoidal compared to the square environment and more impaired performance in the narrow than the broad part of the trapezoid. Indeed, calculating the Bayes Factor to quantify the likelihood of observing differences in positional memory between environments and trapezoid halves based on the decoding errors of the SR grid model compared to a null model of no difference revealed strong evidence for the SR grid model (BF_10_=23.58; we report twice the natural logarithm of the Bayes Factor throughout, see Methods).

In the absence of other positional cues, object locations had to be learned relative to the enclosure boundaries in our task. Can differences in boundary proximity explain the pattern of results in line with more accurate memory for positions near boundaries (Lee et al., 2018)? Due to the specific geometry of the trapezoid, distances of object positions to the closest boundary were smaller in the trapezoid than in the square (t(36)=-10.09, p<0.001, d=-1.66, CI: −2.53; −1.72) and smaller in the narrow compared to broad part of the trapezoid (t(36)=-18.34, p<0.001, d=-3.02, CI: −4.87; −3.92). Hence, the boundary proximity model predicts better memory in the trapezoid and the narrow end of this environment — directly opposite to effects we predicted and observed. Congruent with the beneficial role of boundary-proximity, distances to the closest boundary were negatively correlated with memory scores in the square (Supplemental Figure 4E; t(36)=-4.42, p<0.001, d=-0.73, CI: −0.30; −0.11). In the trapezoid, however, there was no consistent relationship between boundary proximity and memory scores (t(36)=-0.40, p=0.524, d=-0.07, CI: −0.08; 0.05; difference between square and trapezoid: t(36)=-3.45, p<0.001, d=-0.57, CI: −0.09; −0.31); suggesting a differential relationship of boundary proximity and positional memory in non-rectangular environments.

Differences in positional memory in both environments were not due to differential navigation behavior: The excess path length of participants’ navigation paths from the start position of a given trial to the remembered object location did not differ in the trapezoid compared to the square environment or between the two parts of the trapezoid (Supplemental Figure 5AB, square vs. trapezoid: t(36)=-0.95, p=0.144, d=-0.16, CI:-1.02; 0.30; broad vs. narrow trapezoid: t(36)=-0.11, p=0.865, d=-0.02, CI: −0.71; 0.42). Further, walking speeds did not differ between the two environments or the sub-parts of the trapezoid (Supplemental Figure 5CD, trapezoid vs. square: t(36)=-0.01, p=0.973, d=-0.002, CI: −0.08; 0.07; broad vs. narrow trapezoid: t(36)=1.15, p=0.079, d=0.19, CI: −0.02; 0.12). Euclidean distances from the start to the correct object positions were not related to spatial memory performance (Supplemental Figure 5EF, square: t(36)=0.17, p=0.764, d=0.03; CI: −0.05; 0.07, trapezoid: t(36)=0.58, p=0.358, d=0.10, CI: −0.05; 0.09, trapezoid broad: t(36)=1.37, p=0.177, d=0.23, CI: −0.04; 0.18, trapezoid narrow: t(36)=-0.01, p=0.987, d=-0.002, CI: −0.07; 0.08).

To explore participants’ navigation behavior in more detail we next examined their body and head orientation during the replacement period relative to the direction from start to response position in each trial. Both body and head orientation of participants were significantly clustered around this direction in square (body: v=36.84, p<0.001; head: v=36.68, p<0.001) and trapezoid (body: v=36.89, p<0.001; head: v=36.68, p<0.001) and the distributions of mean orientations were not different between the two environments (Supplemental Figure 6AB; body: F(1,72)=0.02, p=0.889; head: F(1,72)=0.14, p=0.709). Similar results were obtained for the body and head orientations when comparing trials targeting objects in the broad and narrow parts of the trapezoid (Supplemental Figure 6CD; body broad: v=36.73, p<0.001; body narrow: v=36.73, p<0.001; difference body: F(1,72)=1.53, p=0.220; head broad: v=36.54, p<0.001; head narrow: v=36.65, p<0.001; difference head F(1,72)=0.05, p=0.830). Hence, we do not expect average facing direction to influence our key comparisons between the two environments or within the trapezoid. The circular variance around each trial’s average body direction did not differ between environments (Supplemental Figure 6E; trapezoid vs. square: t(36)=1.06, p=0.118, d=-0.17, CI: −0.01; 0.04) or the trapezoid parts (Supplemental Figure 6G; narrow vs. broad: t(36)=1.14, p=0.076, d=-0.19, CI: −0.01; 0.05), but the circular variance of participants’ facing directions was greater in the trapezoid than in the square (Supplemental Figure 6F; t(36)=2.57, p<0.001, d=0.42, CI: 0.01; 0.06) and greater in the narrow than the broad part of the trapezoid (Supplemental Figure 6H; t(36)=2.13, p<0.001, d=0.35, CI: 0.004; 0.06). This suggests that participants relied more on visual exploration of the environment, decoupled from body rotations and chosen trajectories in our VR setup. The absence of differences in navigated paths speaks against participants being generally disoriented in the trapezoid or fundamental differences in navigational strategies in both environments, but is in line with our interpretation that the effects we observed go back to distorted positional memory. The object position memory task was designed to probe memory for positions in the broad and narrow part of the trapezoid rather than to evenly sample the environment. Therefore, participants faced more frequently towards the narrow or broad end of the trapezoid when navigating towards remembered positions in the respective part of the environment (Supplemental Figure 7A, broad: v=31.84, p<0.001; narrow: v=28.50, p<0.001). Further, participants’ velocity was higher along the trapezoid’s long-axis (Supplemental Figure 7B, broad: v=27.03, p<0.001; narrow: v=13.51, p=0.001). Do attentional resources and task demands differ between test environments? This seems unlikely as our design included a secondary task in which participants memorized color changes of an extramaze cue and later estimated durations between color change events (see Methods). Mean (t(36)=-0.10, p=0.873 d=-0.02, CI: −33.72; 31.24) and absolute (t(36)=-0.32, p=0.629, d=-0.05, CI: −21.94; 14.67) estimation errors as well as error variability (t(36)=-0.81, p=0.205, d=-0.05, CI: −28.56; 10.24) did not differ between square and trapezoid (Supplemental Figure 10), suggesting comparable attentional resources were available in both environments.

### Mnemonic distortions outside of the environment

Next, we sought to address whether computations based on spatial memories distorted through environmental geometry are systematically biased outside of the learning environment. To this end, we asked participants to estimate distances between the positions of object pairs in two modalities subsequent to the encoding phase in VR (Figure 3AB). In the VR version of the distance estimation task, participants reported distances by walking the respective distance between two remembered object positions in a circular enclosure different from the original square and trapezoidal environments. In the desktop version of this task, they indicated these distances on a subjective scale via a computer mouse (see Methods). Participants successfully completed both versions of the task (Supplemental Figure 8AB; long vs short distances in VR version: t(36)=11.00, p<0.001, d=1.81, CI: 2.83; 4.04; mean±SD of Spearman correlations between true and estimated distances in desktop version: r=0.67±0.19), demonstrating the ability to compute never-experienced distances from pairs of remembered positions. Comparing distances walked in the VR version of the task to true Euclidean distances across all trials revealed an overestimation bias (t(36)=5.78, p<0.001, d=0.95, CI: 1.84; 3.64).

**Figure 3.**
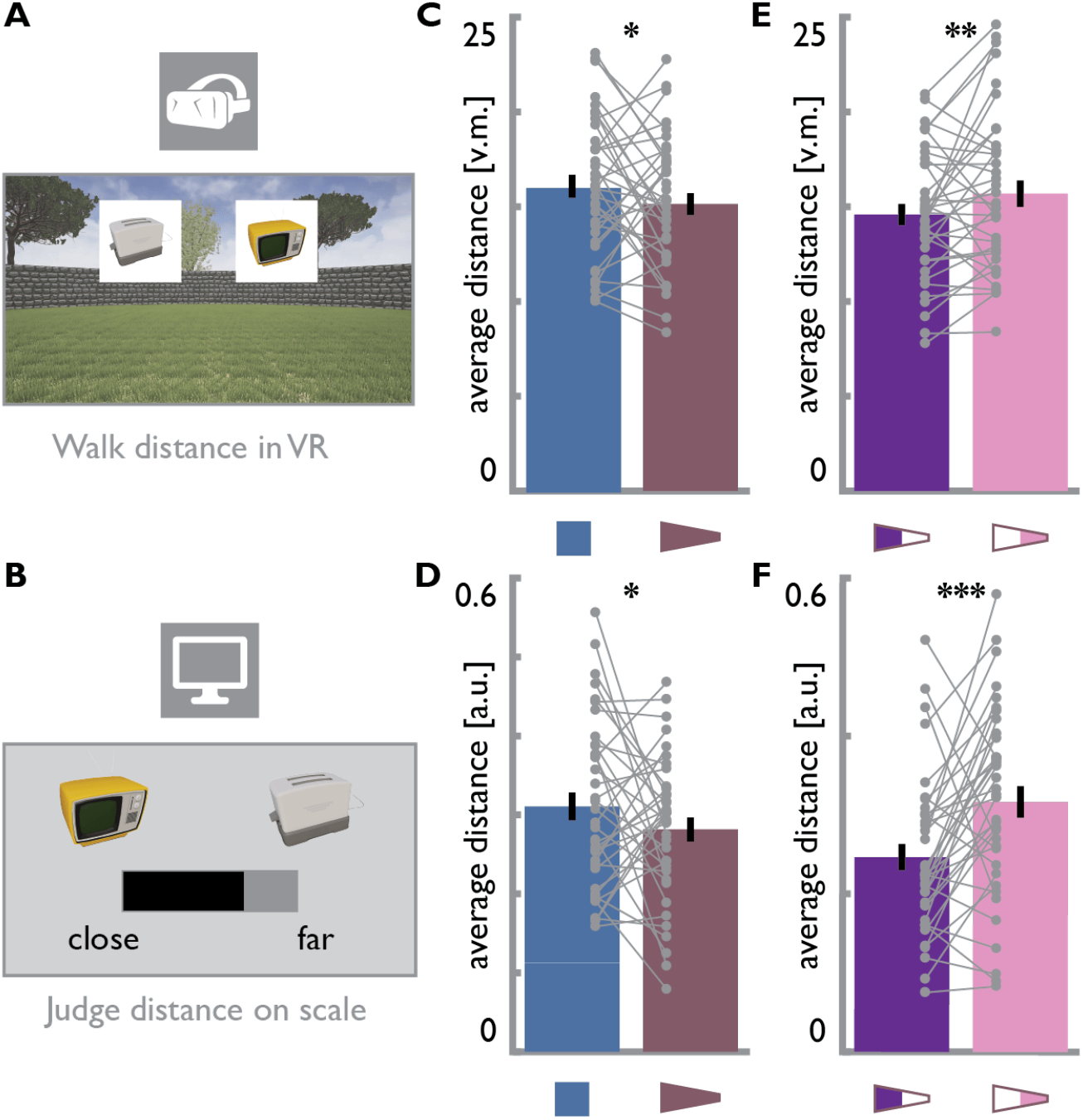
Distortion of distance estimates. **A.** Participants were cued to estimate and walk distances between the remembered positions of object pairs in a circular virtual environment. **B.** In the desktop version of the task, participants adjusted a slider to estimate the distance between object pairs on a subjective scale from close together to far apart. **C, D.** Taking advantage of matched distances between object positions, estimated distances were averaged and compared between environments. Distances were estimated to be shorter in the trapezoid than in the square in both the (**C**) VR and (**D**) desktop version of the task; in line with lower radial frequencies of SR grid patterns in the trapezoid (Figure 1I, Supplemental Figure 3A). **E, F.** Identical distances were estimated to be longer in the narrow than in the broad part of the trapezoid for judgments done in (**E**) VR and (**F**) on a computer screen; consistent with more tightly packed SR grid patterns in the narrow part of the trapezoid (Figure 1J, Supplemental Figure 3B). All bars show mean±SEM and grey circles indicate individual subject data. * p<0.05, ** p<0.01, *** p<0.001

How could distorted grid patterns during encoding bias later distance estimates? The entorhinal grid system is thought to be a central component of the neural substrate supporting vector based navigation — allowing navigational vectors to be calculated by comparing the grid population phases of positions (Fiete et al., 2008; Kubie and Fenton, 2012; Erdem and Hasselmo, 2012; Bush et al., 2015; Banino et al., 2018). In such a system, the difference in grid population phase between locations is expected to be proportional to the Euclidean displacement between them (Bush et al., 2015; Carpenter and Barry, 2016). Thus, in order for a subject to make accurate distance judgements, the relationship between the Euclidean distance and grid phase distance must be held constant in both the presentation and response context. If, for example, a distance was encoded with a population of grid patterns that had been compressed (increased frequency) then attempts to recapitulate that distance with unbiased grid patterns would result in an overestimate in Euclidean space. To determine if successor-based grid patterns were systematically distorted in either the square or trapezoid environment we applied a Fourier approach. Specifically, analysis of the spatial frequency of the SR grid rate maps revealed a sparser packing of grid fields in the trapezoid than the square (Figure 1I, Supplemental Figure 3A, t(98)=3.984, p<0.001, d=0.8, CI: 0.06; 0.18, see methods). Hence, grid phase changed more slowly as a function of distance in the trapezoid, which might result in underestimations of distances relative to the square (Carpenter and Barry, 2016).

Taking advantage of our design in which participants learned a triplet of object positions in each half of an environment with matched inter-object distances, we compared distance estimates between environments. In line with stretched SR grid patterns, distances were judged to be shorter in the trapezoid than in the square in both the VR (Figure 3C, t(36)=1.44, p=0.025, d=0.24, CI: −0.32; 2.20) and the desktop (Figure 3D, t(36)=1.49, p=0.027, d=0.24, CI: −0.004; 0.07) version of the task. This effect was highly reliable between the two versions of the task (Supplemental Figure 8C, Spearman r=0.79, p<0.001, CI: 0.61; 0.88). Next, we tested for a difference between distance estimates for the two parts of the trapezoid. Consistent with higher frequencies of the successor based grid patterns in the narrow compared to the broad part (Figure 1J, Supplemental Figure 3B, t(98)=3.898, p<0.001, d=0.78, CI: 0.06; 0.2), participants estimated the same distances to be longer in the narrow than in the broad part of the trapezoid (VR: Figure 3E, t(36)=2.09, p=0.002, d=0.34, CI: 0.10; 2.15; desktop: Figure 3F, t(36)=3.46, p<0.001, d=0.57, CI: 0.03; 0.11). Again, the difference between remembered distances was highly correlated across the two modalities (Supplemental Figure 8D, Spearman r=0.70, p<0.001, CI: 0.36; 0.89). We calculated surrogate distributions of distance differences between the two halves of the square to contrast the two environments. In both versions of the task, the distance difference of the trapezoid halves differed significantly from the surrogate distributions of distance differences obtained from the square halves (Supplemental Figure 8EF, VR: Z=4.05, p<0.001; desktop: Z=3.68, p<0.001). These findings demonstrate that — across two versions of the task with very different response formats — distance estimates for identical distances are systematically biased in a way consistent with the spatial frequencies of distorted SR grid patterns in the trapezoid. We assessed the likelihood of observing longer distance estimates in the square and in the narrow end of the trapezoid given differences in spatial frequency of the SR grid patterns. Again, the Bayes Factor strongly favored the SR grid model over the null model (VR: BF_10_=15.31; desktop: BF_10_=18.68).

### Reconstruction of remembered locations

What is the structure of deformed memory maps? To reconstruct remembered object positions from estimated inter-object distances, we applied multidimensional scaling to the data obtained in the desktop version of the task (Figure 4A). We extracted coordinates along two dimensions (see Supplemental Figure 9A) which we mapped onto the true coordinates of the trapezoid using Procrustes analysis to match the two configurations of coordinates (Figure 4A and Supplemental Figure 9B, see Methods). We quantified the deviance between the true and reconstructed positions after Procrustes analysis and compared this Procrustes distance to a surrogate distribution of distances obtained from shuffling object-position-assignments to assess the statistical significance of the reconstruction accuracy (Figure 4B). The observed Procrustes distances were significantly lower than the 5th percentiles of the surrogate distributions (Figure 4C, t(36)=-8.48, p<0.001, d=-1.39, CI: −0.21; −0.13), reflecting a close match between true and reconstructed positions. Importantly, re-calculating above-described memory scores with the reconstructed positions led to higher scores compared to the true positions (Figure 4D, t(36)=3.09, p<0.001, d=0.51, CI: 0.0007; 0.003), providing direct evidence that positional memory is used to compute distances between objects and that distorting the spatial map also distorts distance estimates. This effect was also significant when excluding trials targeting objects whose reconstructed position lay outside of the environment (t(36)=1.42, p=0.023, d=0.23, CI: −0.0003; 0.002). The increase in memory scores could be explained if, for each position, reconstructed positions reflect remembered positions in the trapezoid. To quantify this, we calculated the error vectors between the true and remembered positions in the object position memory task and compared these to the error vectors of the reconstructed positions. We observed a strong relationship between the two sets of error vectors as indicated by a significant correlation of their average lengths (Figure 4E, r=0.62, p<0.001, CI: 0.33; 0.79) and a clustering of their orientations (Figure 4F, angular difference of vectors significantly clustered around 0: v=13.77, p=0.001). These findings show that positions reconstructed based on distance estimates were shifted in the same direction as remembered positions and that the magnitude of this shift corresponded to the size of errors in the object position memory task. a

**Figure 4.**
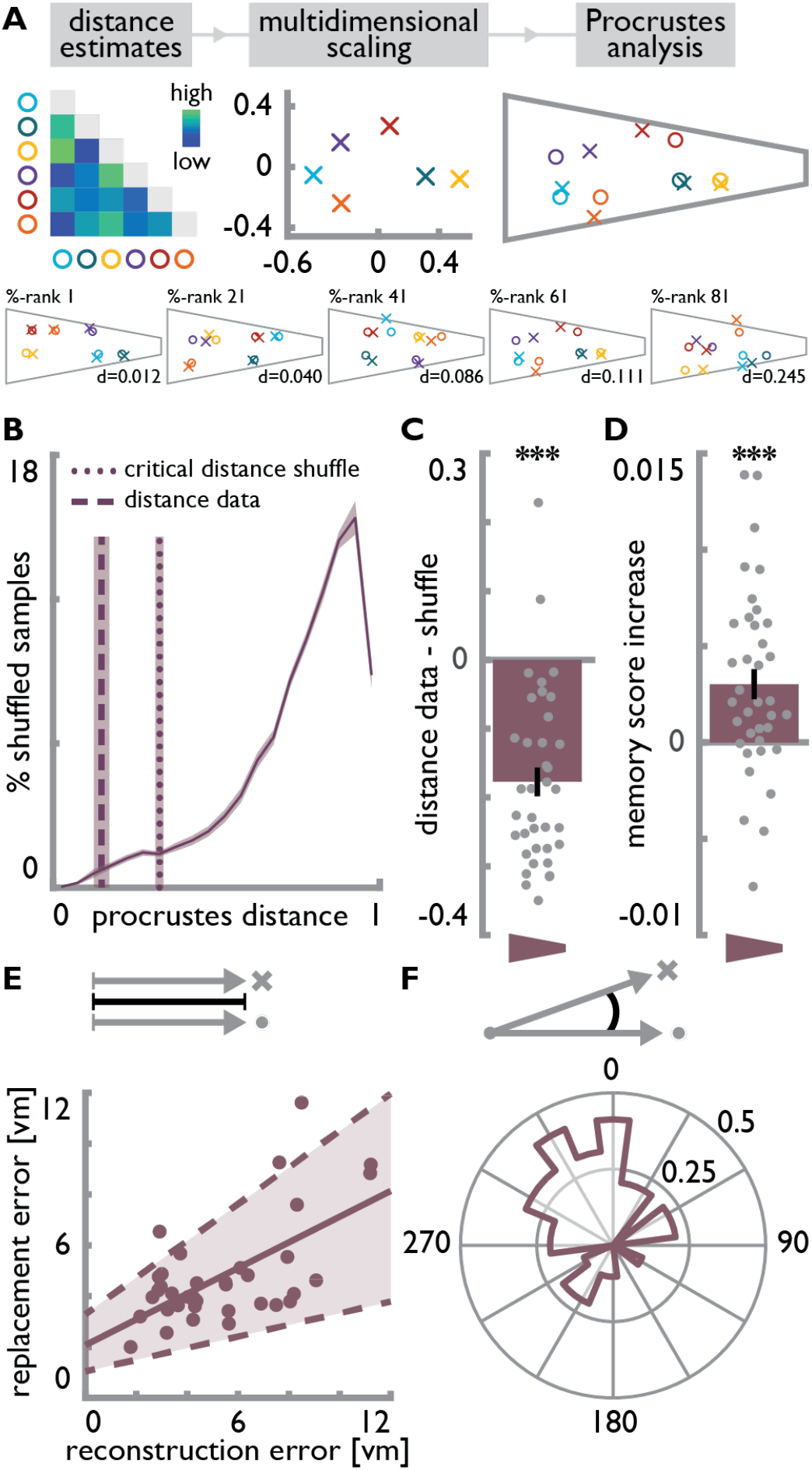
Reconstruction of remembered positions. **A.** Using multidimensional scaling (MDS), coordinates along two underlying dimensions were extracted from pairwise distance estimates. The resulting coordinates were mapped to the original object positions in the trapezoid using Procrustes analysis (see Methods). Color bar indicates estimated distances. Data shown for a randomly selected participant in top panel. Bottom row shows five participants chosen to illustrate reconstruction accuracy across the sample based on the spread of Procrustes distances (data shown for participants at percent ranks 1, 21, 41, 61 and 81 from left to right). Colored circles indicate correct positions and crosses the respective reconstructed positions. **B.** The Procrustes distance quantifies the deviation between true and reconstructed positions as the normalized sum of squared error distances (mean across participants shown by dashed vertical line). For each participant, a surrogate distribution of Procrustes distances was obtained from fitting the coordinates from MDS to coordinate sets with shuffled object identities (solid line). Dotted vertical line indicates the averaged critical Procrustes distance defined as the 5th percentile of the respective surrogate distributions. Shaded areas show SEM across participants. **C.** The Procrustes distances from fitting to true coordinates were significantly smaller than the critical distances of the surrogate distributions. **D.** Memory scores quantifying positional memory within the environment were significantly higher when calculated with respect to the reconstructed rather than the true object positions. **E.** The average error vector lengths from the object position memory task correlate across participants with error vector lengths of the positions reconstructed from distance estimates. Circles show individual participant data; solid line is the least squares line; dashed lines and shaded region highlight bootstrapped confidence intervals. **F.** The angular differences in orientation between the two sets of error vectors cluster around 0°, indicating that errors were shifted in the same direction. Bars in **C** and **D** show mean±SEM and grey circles indicate individual subject data. *** p<0.001

## Discussion

Here, we used immersive VR to demonstrate that environmental geometry can distort human spatial memory. Our data show that positional memory is impaired in a trapezoid compared to a square with the deficits being most pronounced in the narrow end of the trapezoid environment. These findings are consistent with environmentally-induced distortions observed in rodent entorhinal grid-patterns. Equally, they closely mirror predictions drawn from a model grid cell system derived from the eigenvectors of the successor representation (SR). Importantly, mnemonic distortions persisted outside of the environment: Participants estimated identical distances to be different in the square and trapezoid as well as between the narrow and broad part of the trapezoid, underscoring an effect of environmental geometry during encoding on subsequent memory in line with spatial frequencies of SR grid patterns. Moreover, remembered positions reconstructed from these distance estimates directly reflected positional memory during the learning task.

Our findings demonstrate a strong impact of environmental geometry on human spatial memory. We predicted this influence from rodent data in which the six-fold symmetry of grid firing is distorted in a trapezoidal enclosure, with the most pronounced distortions in the narrow part of the enclosure (Krupic et al., 2015). We show degraded positional memory as a function of environmental geometry, in line with larger position decoding errors based on the eigenvector grid patterns of the SR as well as with impaired positional decoding from simulated grid cells with locally distorted firing patterns (Krupic et al., 2018). In concert with evidence for impaired path integration with disrupted grid cell firing in rodents (Gil et al., 2018) and increased path integration errors in older adults with weaker hexadirectional signals measured with fMRI (Stangl et al., 2018), previous studies support the interpretation that the integrity of the grid pattern is beneficial for human spatial memory. The strength of hexadirectional signals and the directional coherence of the orientation of these signals across voxels in the entorhinal cortex are associated with memory performance across participants learning object positions in circular enclosures (Doeller et al., 2010; Kunz et al., 2015). Our findings dovetail with this notion as they demonstrate that environmental geometry, known to compromise grid-patterns in rodents, influences spatial cognition in a within-subject design.

We further demonstrate that distortions persist beyond the encoding environment. The grid cell population phase is thought to provide a mechanism to encode spatial positions and calculate vectors between locations (Bush et al., 2015). As such, distortions of the grid-pattern can decouple the rate of change in population phase from distance in the environment (Carpenter and Barry, 2016). Two positions separated by a given distance will be encoded more similarly when grid patterns are of lower rather than higher spatial frequency. When estimating distances between positions, more similar grid population phases will result in shorter distance estimates (Fiete et al., 2008; Erdem and Hasselmo, 2012; Kubie and Fenton, 2012; Bush et al., 2015; Banino et al., 2018). Consistent with the lower spatial frequencies of SR grid patterns, participants estimated identical distances to be shorter in the trapezoid than in the square. Within the trapezoid, SR grid patterns had a higher spatial frequency in the narrow end and, consistently, participants estimated distances to be greater in the narrow compared to the broad part.

Our results show that human spatial memory was distorted in a trapezoidal environment, suggesting that boundary geometry can distort mnemonic representations. Previous studies have investigated the role of trapezoidal boundary geometry for spatial updating and reorientation. Evidence suggests successful use of trapezoid room geometry for spatial updating in the absence of additional orientation cues. A limited number of symmetry axes was suggested to facilitate the maintenance of one’s orientation in angular in contrast to circular environments (Kelly et al., 2008). Consistently, human participants successfully rely on trapezoidal boundary geometry for heading retrieval (Sturz et al., 2011; Twyman et al., 2018). In our task, wall textures provided additional, unambiguous non-geometric cues for orientation in both environments, making it unlikely that participants were disoriented in either the square or the trapezoid (Hermer and Spelke, 1994; Twyman et al., 2018). This supports our interpretation of the effects reflecting differences in positional memory rather than being driven by disorientation or differences in navigation behavior. Learning positions relative to environmental boundaries recruits the hippocampal formation and is thought to occur incidentally (Doeller and Burgess, 2008; Doeller et al., 2008). Recent evidence suggests that positions near a boundary are remembered more accurately than positions in the center of a rectangular enclosure (Lee et al., 2018). While we replicate the finding that boundary proximity is beneficial for positional memory in the square, this cannot explain the pattern of results we observed. Positions in the trapezoid were closer to the nearest boundary than in the square and, within the trapezoid, positions in the narrow were closer to a boundary than in the broad part. Our findings suggest that boundaries can also distort human spatial memory, in line with grid pattern distortions through environmental geometry.

Prior studies suggested that changing environmental boundaries might influence human spatial cognition in ways consistent with findings from studies of rodent place (O’Keefe and Dostrovsky, 1971) and grid cells (Hafting et al., 2005). Focusing on path integration, one of the core functions assumed for grid cells (Hafting et al., 2005; McNaughton et al., 2006; Moser et al., 2017), biases in human navigation have been reported to follow predictions derived from grid cell firing (Chen et al., 2015). In particular, the experimental design in Chen et al. (2015) built upon the observation that rodent grid-patterns rescale to match changes made to the geometry of already familiar enclosures (Barry et al., 2007). Expansions and compressions of boundaries relative to preceding trials resulted in under- and overshoots of the return path in a path integration task, when the path included a component along the manipulated boundary dimension (Chen et al., 2015). This illustrates how, through environmental change, altering the rate of change in grid cell population phase in relation to distance traveled can introduce biases in human navigation (Carpenter and Barry, 2016; Chen et al., 2015). As described above, translating this idea to the memory-based estimation of distances between locations might explain the diverging judgments of identical distances observed in our data. Expansions and compressions of virtual environments have further been demonstrated to impact spatial memory in humans and, under conditions of environmental change, positional memory follows models of place cells and boundary proximity (Hartley et al., 2004; Schuck et al., 2015). While the studies described above indicate how boundary manipulations in familiar environments influence spatial behavior, we built upon work showing that distorted grid-patterns persist in static trapezoid environments even with prolonged experience (Krupic et al., 2015). Our findings suggest that distortions of the brain’s spatial metric can result in mnemonic distortions under constant boundary conditions within a specific environment and even outside of this encoding environment.

We opted for a purely behavioral experiment; our hypotheses, experimental design and analysis however directly built upon findings from electrophysiological recordings of grid cells in rodents (Krupic et al., 2015). We employed highly immersive VR technology to enhance the impact of environmental geometry on spatial cognition and engage proprioceptive, vestibular and motor systems during the task. Currently, immersive VR does not allow the concurrent recording of neural data. The contribution of locomotor cues to the experience of navigation in general has been emphasized previously (Taube et al., 2013) and recent studies in rodents have used gain manipulations in VR to emphasize the contributions of locomotor cues to grid cell firing specifically (Campbell et al., 2018; Chen et al., 2019). Having established the impact of environmental geometry on human spatial cognition, an exciting question for future research would be to combine manipulations of environmental geometry with neuroimaging techniques such as fMRI to study the deformations of the cognitive map we describe here in the brain. To do so, an important measure could be the hexadirectional signal that can be observed in the human entorhinal cortex (Doeller et al., 2010). Beyond fMRI, an exciting future avenue is paved by the development of new magnetoencephalography systems, which might allow the combination of immersive VR with recordings of neural data (Boto et al., 2018).

As large parts of human indoor navigation take place in rectangular rooms, the novelty of a trapezoidal enclosure in our task might be considered as a factor contributing to impaired performance compared to the square. Such an effect of unfamiliarity with polarized environments, however, would not predict the observed within-environment differences in performance. Further, participants’ walking speeds did not differ between environments and their paths from the start to the remembered object position were not more or less direct. Thus, none of our control measures provided evidence for fundamental differences in navigational performance between the environments per se. Additionally, the detection and encoding of color change events was not affected by the environmental manipulation, speaking against an effect of increased task demand in the trapezoid as sufficient attentional resources were available for this secondary task.

Importantly, the effects we observed in positional memory persisted outside of the environment as demonstrated by the differential estimates for matched distances between positions within the different parts of the trapezoid. These distortions were highly reliable across response modalities, demonstrating a general, task-invariant mnemonic effect. Our findings dovetail with asymmetric distance judgments between landmarks and non-landmarks as well as overestimations of distances as a function of intermediary boundaries (Cadwallader, 1979; Sadalla et al., 1980; Thorndyke, 1981; McNamara, 1986; McNamara and Diwadkar, 1997; Newcombe et al., 1999). Beyond boundaries separating positions, our findings demonstrate that distance estimates can be influenced through the geometric arrangement of boundaries. The response profiles observed in the VR version of the task revealed a general tendency to overestimate distances between positions, consistent with previous studies reporting overestimations of navigated distances (Brunec et al., 2017) and spatial scale in map drawings (Jafarpour and Spiers, 2017). We used the distances estimated on a subjective scale in the desktop version of the task to reconstruct remembered positions. Accounting for the distortions in participants’ memory by using these reconstructed positions to re-compute memory scores yielded increased performance scores. This illustrates the close match between positions reconstructed from distance estimates and positional memory within the environment, and demonstrates that, consistent with the formation of cognitive maps (O’Keefe and Nadel, 1978), distances never directly experienced in the task were computed from remembered positions. Grid cells have been suggested to support this kind of vector computation (Bush et al., 2015; Banino et al., 2018). This is further in line with evidence for the involvement of the entorhinal grid system in imagination (Bellmund et al., 2016; Horner et al., 2016) and theoretical accounts proposing a role for spatially tuned cells in memory (Byrne et al., 2007; Buckner, 2010; Hasselmo, 2011).

Environmental geometry systematically biased memory-based computations outside of the trapezoid environment, thereby linking our findings to a growing body of literature implicating grid-cell computations in cognitive functions beyond navigation (Bellmund et al., 2018a). For example, grid-like hexadirectional signals were also observed during trajectories through an abstract space spanned by the dimensions of neck and leg length of stick figure birds (Constantinescu et al., 2016). Collectively, these findings point towards a role of the entorhinal grid system in mapping cognitive spaces (Bellmund et al., 2018a). As proposed for navigable space (Hafting et al., 2005; McNaughton et al., 2006; Bush et al., 2015; Moser et al., 2017; Herz et al., 2017), the regular firing patterns of grid cells might provide a metric for these spaces allowing the efficient encoding of specific stimuli located at different positions within a space. Speculatively, correlated feature dimensions or feature spaces in which subsets of feature combinations are impossible might distort how grid cells map these spaces in a similar way as environmental geometry distorts grid cell firing patterns, resulting in biased representations similar to the distortions of spatial memory observed in this study.

In conclusion, our data show distortions of human spatial memory consistent with the changes induced in rodent grid cell activity by the geometry of highly polarized enclosures. These distortions persist outside of the environment, indicating an enduring impact of environmental geometry on memory. In line with the proposed roles for grid cells in navigation and mapping feature dimensions beyond navigable space, these findings suggest that environmental geometry might be able to distort the metric of cognitive representations.

## Methods

### Participants

53 Participants between the age of 18 and 30 were recruited from the Norwegian University of Science and Technology. All participants provided written informed consent before participation, and all research procedures were approved by the regional ethics committee (REC North, reference number 2017/153). Sample size was based on a power calculation assuming a small to medium effect (d=0.4) of environmental geometry on human spatial cognition, resulting in a sample size of 52 to achieve a statistical power of 80% (∝=0.05, two-tailed test). 39 participants (mean age 23.8±2.5 years, 36% female) completed the experiment (14 incomplete datasets due to technical difficulties with the VR setup or motion sickness). Two participants were excluded due to poor memory performance defined as average replacement errors more than 1.5 times the interquartile range larger than the upper quartile of average errors in the sample. Thus, 37 participants entered the analyses.

### Overview

We designed our experiment to test distortions of spatial memory as a function of environmental geometry. Figure 1A provides an overview of the experiment structure. Participants were first (Figure 1A: I) familiarized with the VR setup before beginning the object position memory task in the first environment. The object position memory task (Figure 1A: II & IV) was carried out in a trapezoidal or square environment for 20 minutes each, with the order of environments counterbalanced across participants. Subsequent to navigating an environment, participants were prompted to estimate the durations between occasional color change events encountered in that environment (Figure 1A: III & V). In the last two tasks, participants were asked to estimate distances between pairs of objects in VR and on a computer screen (Figure 1A: VI & VII), respectively. The design of each task and the corresponding analyses are described in detail in the following sections. All analyses were performed using Matlab (Release 2017a, The MathWorks, Inc.) and statistical tests (two-tailed unless stated otherwise, alpha-level 0.05) were performed using resampling procedures as implemented in EEGLAB (Delorme and Makeig, 2004). Specifically, test statistics were compared against a surrogate distribution obtained from 10000 bootstrap samples respecting within-subject dependencies. We report 95% confidence intervals of mean differences; bootstrapped using the bias-corrected and accelerated percentile method as implemented in the Matlab function ‘bootci’. Bootstrapped p-values and confidence intervals do not rely on Student’s t-distribution and therefore 95% confidence intervals can include 0 even if the p-value is smaller than 0.05. Cohen’s d was calculated by dividing the t-value by the square root of the number of participants (Lakens, 2013). Circular statistics were implemented using the Matlab-based Circular Statistics Toolbox (Berens, 2009).

### Virtual reality

Aiming to maximize the feeling of immersion and thereby the impact of environmental features we employed state of the art VR technology consisting of a head mounted display (HMD, Oculus Rift CV1) and a motion platform (Cyberith Virtualizer). Participants wore low-friction overshoes and were strapped into a harness attached to the motion platform’s ring system allowing free rotations. To navigate the virtual environments, participants were instructed to lean slightly into the ring construction to slide the front foot backwards across the sensors of the low-friction base plate of the motion platform while taking a step forward with the back foot (see Supplementary Video 1), generating translational movement in the current forward direction determined by the orientation of the participant in the ring system (Cakmak and Hager, 2014). Head movements were tracked in 3D using the HMD’s tracking system and the virtual environments were displayed to both eyes separately at a resolution of 1080 x 1200 pixels and a refresh rate of 90 Hz. The virtual environments were created and presented using the Unreal Engine (v.4.13.2, Epic Games Inc., 2017) and participants’ eye height was set to 1.80 virtual meters (vm). Participants were familiarized with the VR setup in a circular environment (45.74vm in diameter) consisting of a grass floor curtailed by a wall (height 3.75vm). A set of trees spread around the outside the environment served as cues for orientation. During familiarization, participants practiced walking and turning by navigating the circular environment to collect coins appearing at random positions in the environment. Participants were instructed to walk towards the coins and collect them via button presses on a handheld controller. Additionally, this familiarization period served as a practice for the time estimation task (see below).

### Object position memory task

Participants performed an object position memory task during which they iteratively learned the positions of six objects in a trapezoidal environment (36vm×76vm×8vm×76vm) with side lengths proportional to the enclosure rodents explored in a study reporting distortions of grid cell firing patterns (Krupic et al., 2015). To establish a behavioral baseline, participants performed this task also in a square control environment (40.27vm×40.27vm) with equal surface area. There were no distal cues outside of the environment to enforce spatial learning based on environmental geometry. To facilitate orientation, each wall was presented in a unique color. Both environments had a grass floor and a blue sky with moving clouds was visible (Figure 1B). Participants performed the task for 20 minutes in each environment with the order counterbalanced across participants. In each environment, participants learned the positions of six everyday objects presented as three-dimensional models. The assignment of objects to arenas and positions was randomized across participants.

In each trial of an initial learning phase, participants navigated to a start position indicated by a traffic cone. Then, an object was shown at its predefined position in the environment and participants were instructed to navigate to the object, collect it via button press and memorize its position. Each object was shown once and the order of objects was randomized. In the subsequent test phase (Figure 1B), participants again navigated to start positions. Upon arrival, a picture of one of the objects was shown as a cue for 3 seconds in front of the participant, prompting participants to navigate to where they remembered this object in the environment. Participants indicated the remembered position via button press after arrival and received feedback about their accuracy in the form of one of five smiley faces. The object then appeared at its correct position and participants collected it before the beginning of the next trial. Participants completed 30.54±6.71 and 30.38±8.09 (mean±SD) test trials in square and trapezoid, respectively (number of trials not significantly different: t(36)=0.18, p=0.759).

The order of trials was randomized for mini-blocks of six trials, so that within a mini-block each object was sampled once and no two consecutive trials sampled the same objects. A triplet of object positions (Figure 1C) was randomly generated for each participant with a minimum distance of 11vm between object positions and a minimum of at least 3vm to the nearest boundary. Positions were constrained so that the connection between two objects was parallel to the long-axis of the trapezoid or one of the walls of the square. The third object was placed at an angle ranging from 90°-120° relative to the first two with the same distance to one of the objects as between the first two. Such a triplet of positions was placed in both the narrow and broad part of the trapezoid defined based on the midpoint of its long-axis and the left and right part of the square. Placing triplets of objects with matched distances in each part of the environment allowed direct comparisons of remembered distances between environments and their sub-parts (see distance estimation tasks). Since cues were only shown once participants arrived at the start position of a given trial, participants never walked the direct path between two objects. Distances from start to target object positions (mean and standard deviation square: 18.66±4.65vm; trapezoid: 19.92±8.50vm; trapezoid broad: 21.10±10.95vm; trapezoid narrow: 18.73±4.67vm) did not influence spatial memory performance (Supplemental Figure 5EF).

### Positional memory

Raw positional memory errors were quantified as the Euclidean distance between the correct position of an object in the environment and the position remembered by the participant. To limit the influence of outlier trials we excluded trials with errors larger than 1.5 times the interquartile distance above the upper quartile of errors for each participant (mean±SEM number of trials excluded =3.35±0.26) from all further analyses. Average positional memory errors were compared across environments using a bootstrap-based paired t-test (Figure 2A). To account for the fact that despite equal area larger errors are possible in the trapezoid compared to the square control environment, we subsequently quantified performance using memory scores. Specifically, we generated a distribution of 1000 random locations uniformly covering each environment and quantified for each trial the proportion of locations further away from the correct object position than the position indicated by the participant. Importantly, calculating memory scores based on the distribution of possible errors for each target position yields a measure comparable across positions and environments (Jacobs et al., 2016) with a chance level of 0.5 for random performance and scores closer to 1 for high performance. To test the hypothesis of degraded spatial memory in the trapezoid memory scores were compared across environments using a bootstrap-based paired t-test (Figure 2B).

In a next step, we aimed to test the more specific hypothesis of increased degradation of positional memory in the narrow compared to the broad part of the trapezoid derived from the larger distortions of firing patterns of grid cells in this part of the environment (Krupic et al., 2015). We used a bootstrap-based t-test to test whether positional memory errors differed between the narrow and broad part of the trapezoid (Figure 2D). Outlier participants were excluded based on our standard criterion of values more than 1.5 times the interquartile range above or below the upper or lower quartile, respectively (see Supplemental Figure 4B for full dataset). Distributions of possible errors can differ also for positions within the same environment. Therefore, we also tested whether memory scores differed between the two parts of the trapezoid (Figure 2E).

Since the rotationally symmetric geometry of the square does not pre-define how to calculate the difference in positional memory, we created a surrogate distribution by shuffling which half of the environment was to serve as the subtrahend and minuend for the error difference across participants. For each permutation, we calculated the error difference for objects located in the two halves of the square. The positional memory error difference observed in the trapezoid was smaller than the 5^th^ percentile (one-tailed test) of the surrogate distribution obtained from 10000 permutations, (Supplemental Figure 4C). The shape of the surrogate distribution did not differ from normality (Kolmogorov-Smirnov test, D=0.01, p=0.277), we hence used it to convert the p-value reflecting the number of occurrences of smaller memory ratios in the surrogate distribution into a Z-statistic. To visualize response behavior in the two parts of the trapezoid we collapsed across all trials from all participants for objects located in the broad and narrow part of the arena. Response positions were centered on the respective true positions and divided into 50×50 square bins with a side length of 0.6vm. The resulting histogram was smoothed using a Gaussian kernel with a standard deviation of 0.5vm and plotted as a heatmap (Supplemental Figure 4D). To test the influence of the distance to the nearest boundary on positional memory we calculated the Pearson correlation between the Euclidean distance to the closest boundary and the memory scores across all trials from an environment for each participant. We tested the resulting correlation coefficients against 0 and between the environments using bootstrap-based t-tests. Negative correlation coefficients indicate better memory closer to the boundary.

### Parameters of navigation

To assess whether differences in navigation behavior might underlie the observed differences in positional memory, we analyzed navigational performance in the replacement phase of each trial, where participants navigated to the remembered position of a cued object. For each trial, we calculated the Euclidean distance between the start position and the response location and subtracted it from the length of the path walked by the participant. This excess path length measures the directness of the paths taken, potentially reflecting the degree of certainty about the trajectory as increased uncertainty might lead to more turns and longer paths. We contrasted averaged excess path lengths between the two environments and the broad and narrow part of the trapezoid (Supplemental Figure 5AB). Likewise, we contrasted average walking speeds during the replacement phase between the environments and trials targeting objects from the two trapezoid parts (Supplemental Figure 5CD).

Further, we assessed whether the distance from a trial’s start position was related to the accuracy of object position memory in a consistent way across subjects. For each subject, we calculated the Spearman correlation coefficient between the distances from start to true object positions and positional memory as defined by the Euclidean distances between true and remembered object positions. The resulting coefficients were tested against 0 for all trials in the two environments separately (Supplemental Figure 5E) or for trials probing objects in the narrow and broad part of the trapezoid, respectively (Supplemental Figure 5F).

In a next step, we assessed rotations participants made during the replacement phase of the trial. To this end, we centered the rotation of the body as measured by the orientation of the motion platform’s ring construction and the orientation of the participant’s head as tracked by the HMD on the direction from start to response position. We averaged orientation values for trials within square and trapezoid or broad and narrow part of the trapezoid, respectively, and tested for clustering around 0° using V-tests and differences of averaged orientation values between conditions using Watson-Williams tests (Berens, 2009) (Supplemental Figure 6, top row). Additionally, we quantified the circular variance of centered orientation values and contrasted it across conditions (Supplemental Figure 6, bottom row). None of these measures suggested influences of navigation behavior per se on the key conclusions of the paper.

Further, we tested the sampling of directions separately for trials targeting objects in the narrow and broad part of the trapezoid. For each of 36 angular bins with a width of 10° we computed the proportion of time points for which participants’ bodies faced the direction of that bin. We averaged these proportions across participants for the polar histogram in Supplemental Figure 7A. To test whether angular sampling was biased towards the long and short base of the trapezoid we calculated the angular mean for each participant and used v-tests to test for a clustering around 180° and 0°, respectively. Next, we quantified average movement velocity for each direction bin (Supplemental Figure 7B). We weighted directions by average velocity to compute a circular mean for each participant. Again, we tested using v-tests whether the resulting circular means clustered around 180° and 0° for trials where target objects were located in the broad and narrow part of the environment, respectively.

### Distance estimation tasks

After completing the time estimation task following the second object position memory task in the second environment, participants estimated distances between pairs of object positions in two modalities: on a computer screen and by walking the actual distances in VR.

#### Virtual reality

Participants were placed in the same circular virtual arena as during the familiarization session. Each trial began with an arrow pointing to the middle of the arena, with the arrow appearing at a random location on the arena floor. After participants positioned themselves on the base of the arrow, images of two objects were presented in front of them for 3 seconds (Figure 3A). Participants were instructed to walk the distance they remembered the objects to be apart based on the object position memory task while following the direction indicated by the arrow. When participants terminated a trial via button press, a checkmark was presented to indicate the successful registration of the response and the next trial began. Due to time constraints this task was restricted to distances between objects within a triplet, resulting in 12 trials making up a block. Trial order within blocks was randomized with the constraint that trials with objects from the two environments alternated. Participants completed two blocks with a short break in between.

Since only distances within a triplet of positions were tested in this task, participants’ averaged estimates for the long and short distances were compared using a bootstrap-based paired t-test as an indicator of successful task performance (Supplemental Figure 8A). To test whether distance estimates for the same distances differed between environments or the narrow and broad part of the trapezoid, we took advantage of the fact that true distances were matched across position triplets and thereby environment parts. Distance estimates for the two triplets within an environment were averaged and contrasted between the square and trapezoid using a bootstrap-based t-test (Figure 3C). Similarly, response distances within a triplet were averaged and compared between the narrow and the broad part using a bootstrap-based t-test (Figure 3E). As for the difference in positional memory, we created a surrogate distribution to compare the distance estimation difference observed between the trapezoid halves to the square by shuffling across participants which half of the square was to serve as the minuend and subtrahend for the distance difference in each of 10000 permutations. The distance difference observed in the trapezoid was more extreme than the 2.5^th^ and 97.5^th^ percentiles (two-tailed test) of this surrogate distribution (Supplemental Figure 8E). The shape of the surrogate distribution did not differ from normality (Kolmogorov-Smirnov test, D=0.01, p=0.200).

#### Computer monitor

Afterwards, participants were instructed to estimate distances between object pairs on a desktop computer setup. Images of objects on a white background, as well as an adjustable horizontal bar with the labels ‘close together’ on the left and ‘far apart’ on the right were presented on a computer screen (Figure 3C). Again, participants were instructed to estimate how far objects were apart during the object location memory task. Here, they indicated their response by adjusting the horizontal bar with a computer mouse, after which a grey screen was shown for 500 milliseconds. All possible combinations of distances were probed, i.e. also comparisons across triplets, yielding subjective distances between all pairs of object positions in an environment. Each of the 15 combinations of object pairs per environment was probed twice, resulting in a total of 60 trials. Trial order was randomized with the constraint that each possible pair of objects was sampled before any object combination was sampled for the second time. Each object was shown once on the left and once on the right side of the screen in the two trials sampling a given object pair. This distance estimation task as well as the time estimation task was presented using the Psychophysics Toolbox (Brainard, 1997) for Matlab (Release 2016a). General performance in this task was assessed by calculating Spearman correlations between the estimated distances and the respective true distances (Supplemental Figure 8B). Further, the distance estimates were contrasted between environments and between the narrow and broad part of the trapezoid in the same way as described above. The surrogate distribution obtained for comparison to the square did not differ from normality (Kolmogorov-Smirnov test, D=0.01, p=0.167).

### Reconstructing remembered positions

To reconstruct remembered object positions in the trapezoid from distance estimates, multidimensional scaling (MDS) was applied to the distance estimates obtained in the desktop version of the task as only here distances between all pairs of positions were estimated. Estimated distances were normalized to a range from 0 to 1 and averaged across the two repetitions of each object pair and subjected to MDS to recover coordinates reflecting this distance structure using metric stress as the cost function and a random initial configuration of points. Our approach assumes that two dimensions underlie the object location memory formed during the navigation task. To assess whether this assumption holds, we compared the model deviance of general linear models predicting the distances between true positions from the positions recovered from MDS for different numbers of dimensions. As expected, unexplained variance was substantially decreased when using two instead of one dimension, but no clear improvement resulted from a larger number of dimensions (Supplemental Figure 9).

To match the coordinates resulting from MDS to the original positions in the virtual environment we used Procrustes analysis allowing translation, scaling, reflection and rotation (see Bellmund et al. (2018b) for an application of the combination of multidimensional scaling and Procrustes analysis to fMRI data). The goodness of fit, the Procrustes distance, was quantified by the normalized sum of squared errors between reconstructed and true coordinates and was compared to Procrustes distances resulting from Procrustes analyses of the MDS coordinates and sets of coordinates in which the assignment of object identity to position was shuffled, yielding a surrogate distribution from all 720 possible permutations. Specifically, we tested on the group level whether the fits between reconstructed coordinates and true coordinates were better than the fits constituting the 5^th^ percentile (reflecting the threshold for statistical significance at an alpha-level of 0.05) of each participant’s surrogate distribution (Figure 4BC). The reconstructed coordinates are visualized as heatmaps in Supplemental Figure 9B following the same procedure as described above.

To test whether the reconstructed positions indeed reflected participants’ memory in the object position memory task, we re-calculated the memory scores as described above but with the coordinates resulting from the Procrustes analysis instead of the true object positions as goal positions (Figure 4D). To rule out that objects whose positions were reconstructed to be remembered outside of the environment were driving the effect, we excluded all affected trials from the memory score calculation in an additional control analysis. To describe the overlap between positions reconstructed from distance estimates and performance in the object position memory task, we calculated error vectors based on the true object positions for both the reconstructed positions and the response positions from the object position memory task. Specifically, we tested whether error vectors were of a similar length and had a similar orientation to demonstrate that positions were shifted by a similar distance and in a similar direction. We quantified the match between average error vectors of response and reconstructed positions by correlating their lengths using Pearson correlation (Figure 4E). We further probed these error vectors’ similarity in orientation by averaging the angular differences between vectors from the correct to the respective response and reconstructed positions for each participant and testing the resulting circular means for a clustering around 0° using a V-test (Figure 4F).

### Time estimation task

To probe whether attentional demands differed between environments we included a secondary task while participants performed the object position memory task. If attentional demand differed across environments, we would expect to see differences in the secondary task performance. In the sky above each arena a ring was presented, which changed color four times during the object position memory task per environment. The ring remained in a given color for an interval between 2 and 6 minutes and participants indicated color changes via button presses and were instructed to remember the order of colors and the duration for which each color was presented. While different colors were presented in the two environments, the intervals between color changes were constant across environments allowing for a comparison of temporal memory between square and trapezoid.

After completing the object position memory task in an environment, participants were placed in front of a computer screen to estimate the time between color changes before continuing with the next part of the experiment (Figure 1A). On a white screen, two pairs of consecutive colors were shown and participants indicated the time interval they remembered to separate the two color changes in minutes and seconds, e.g. how much time passed between the ring changing color from blue to yellow and changing from yellow to green. Participants were cued to estimate the time between all six possible combinations of color changes per environment. To ensure full understanding of this task, participants estimated intervals between color changes occurring at random times between every 30 and 120 seconds during the familiarization phase prior to the object position memory task. Overall performance in this task was quantified using Spearman correlations between the correct and estimated time intervals before specifically comparing average estimation errors, absolute estimation errors and the standard deviation of estimation errors across environments (Supplemental Figure 10).

### Successor representation grid patterns

Following Stachenfeld et al. (2017), simulated grid cells were generated using the first 50 non-constant eigenvectors of the successor representation (SR) matrix under a uniform random walk policy for a square (90×90cm) and a trapezoid (length 187cm, parallel walls 90cm and 20cm, based on Krupic et al. (2015)). Eigenvectors were thresholded at zero and scaled to a peak firing rate of 30Hz. To assess the distortion of SR grid fields within the trapezoidal environment, we divided the square and trapezoid environments into halves across the longest axis and computed the spatial autocorrelation of the SR grid rate maps on each half. Grid similarity was then calculated by taking the Pearson correlation between the autocorrelations from each half of the environment (Figure 1F). Since the shortest dimension of the trapezoid was 20 spatial bins, a circular window of radius 40 spatial bins was used to compare the autocorrelations ensuring the window for comparison never went off the autocorrelation map.

The decoding analysis sampled each spatial bin (1 cm square) of the environment 30 times. On each occasion, for each of the 50 cells, spikes were randomly generated for a 100 ms time window according to a Poisson process with rate parameters equal to the SR grids’ firing rates in that bin. Locations were then decoded using the maximum likelihood estimate of all 50 SR grid spike counts and errors were calculated as the Euclidean distance between the true and decoded locations (Figure 1GH) (Towse et al., 2014; Stemmler et al., 2015). Decoding errors were normalized so that errors in the square and broad part of the trapezoid had a mean of 1 for the comparison between environments and trapezoid halves, respectively.

To analyze differences in the spatial frequencies of the SR grids, we calculated the 2D fast Fourier transform (FFT) for each rate map and reparametrized the FFT into polar coordinates. Power spectra were then considered solely as a function of radial frequency by averaging across the angular component of the FFT (Supplemental Figure 3). Finally, we calculated the mean radial frequency of each SR grid and compared these across and within environments (Figure 1IJ). Statistical significance of all SR grid pattern analyses was assessed using standard two-sample t-tests with the effect size Cohen’s d being the difference between means divided by the pooled standard deviation.

Likelihood analyses were carried out separately for both the positional memory and distance estimation tasks. For each task, the proportional differences in participants’ responses from the square-trapezoid and within-trapezoid contrasts were compared to the distribution of proportional differences expected by the SR model. The same was done for a null model using the same distribution of proportional differences as the SR, but with a shifted mean to predict no overall difference in the square-trapezoid and within-trapezoid contrasts. Likelihoods from the square-trapezoid and within-trapezoid contrasts were combined within tasks to give the likelihood of the human responses given each of the models for both the positional memory and distance estimation tasks. The Bayes factor B_10_ was calculated as the ratio between the two model likelihoods. We report twice the natural logarithm of the Bayes Factor (2ln(B_10_)) as it has a similar scale to familiar likelihood ratio test statistics (Kass and Raftery, 1995). According to the conventions by Kass and Raftery (1995), 2ln(BF_10_)>10 constitutes very strong evidence for the alternative over the null model.

## Supplemental Figures

**Supplemental Figure 1.**
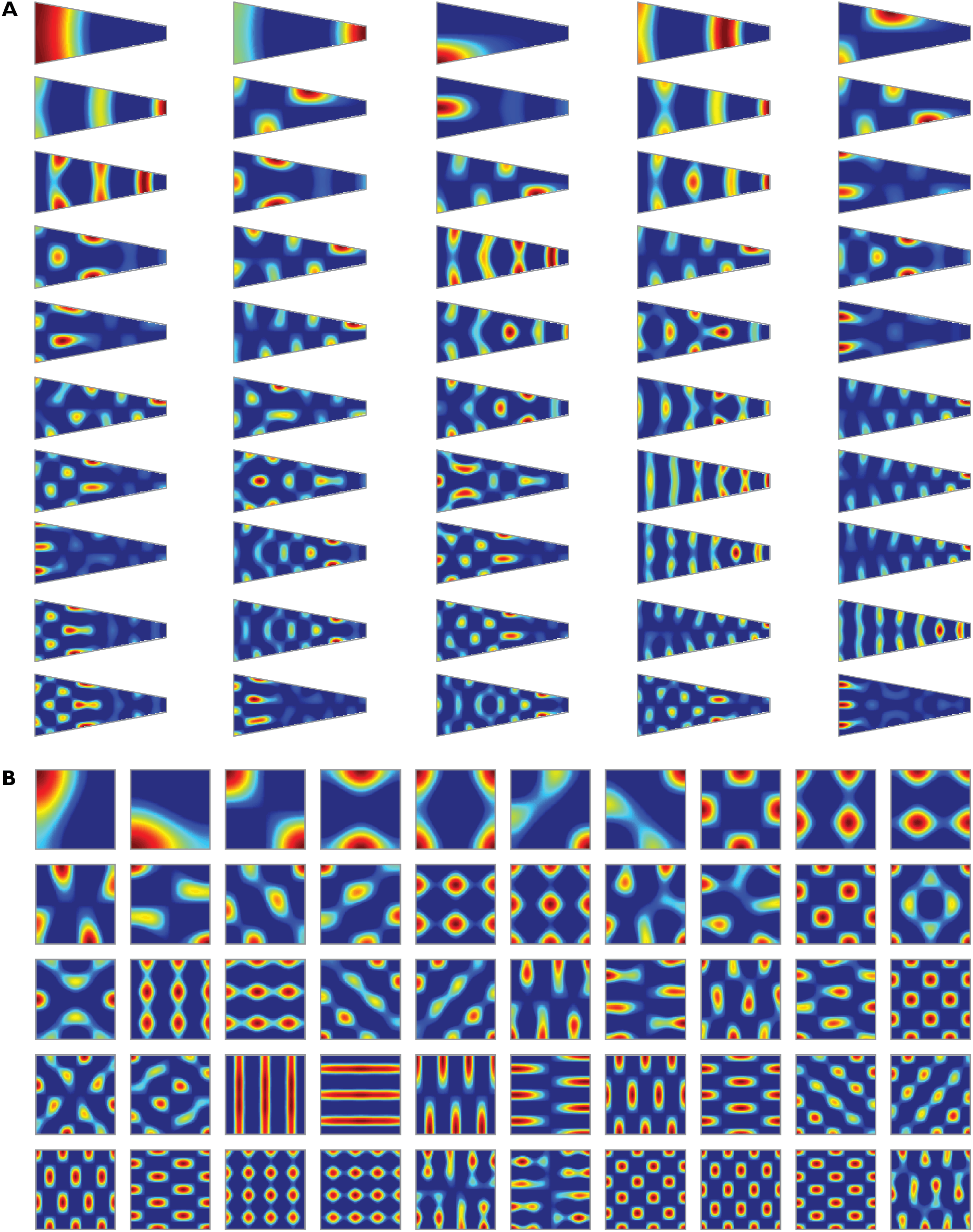
Successor representation eigenvectors. **A, B.** The first 50 eigenvectors of the successor representation from a trapezoid and a square environment were used for analysis.

**Supplemental Figure 2.**
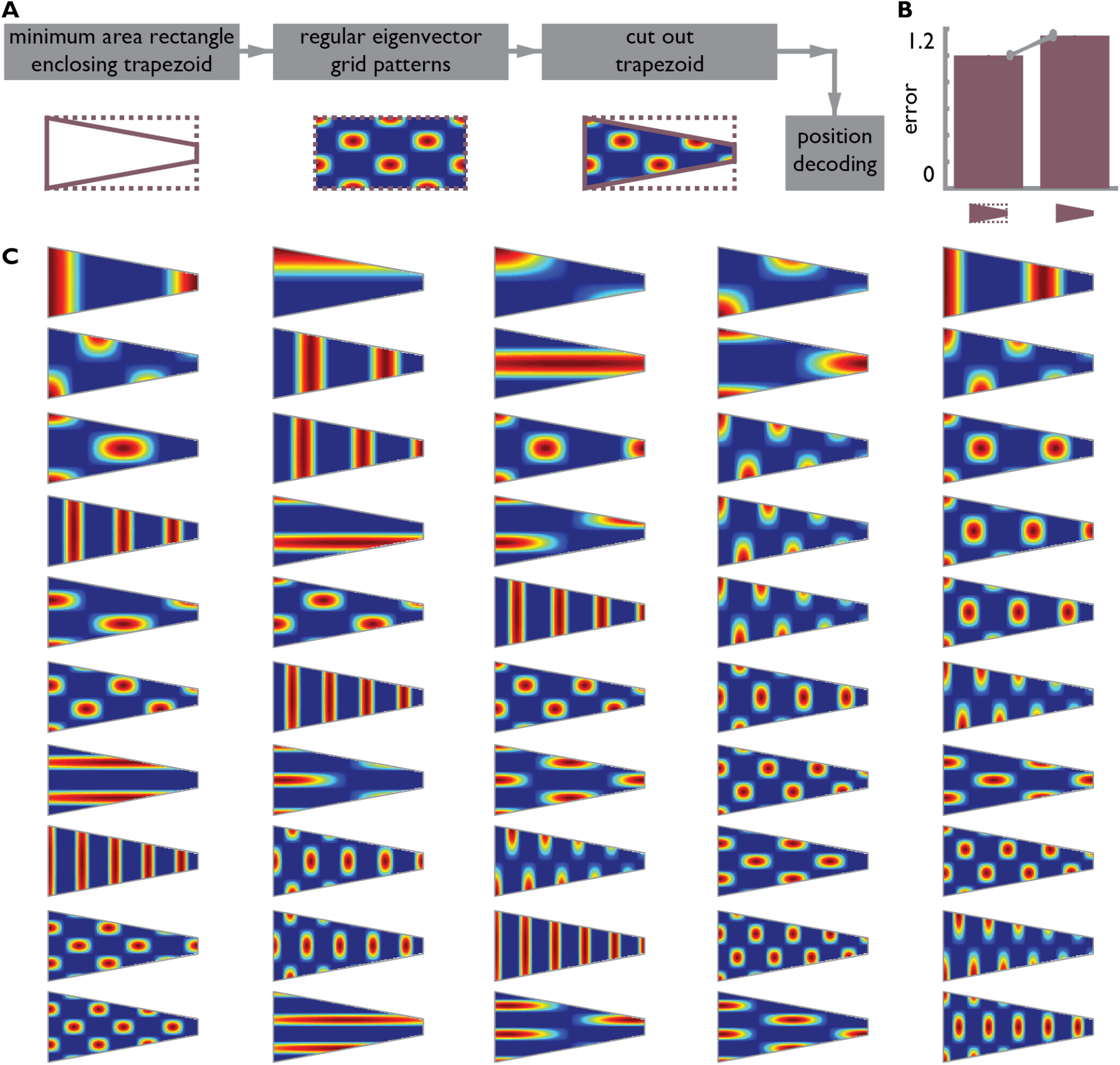
Position decoding based on the smallest rectangle enclosing the trapezoid. **A.** To demonstrate that worse position decoding in the trapezoid is due to the distorted eigenvector grid patterns and not the elongated shape of the environment we analyzed the eigenvectors of the smallest rectangle enclosing the trapezoid. We repeated the position decoding on the area of the trapezoid based on the SR grid patterns from the smallest rectangle. **B.** Position decoding errors were larger when the analysis was based on the distorted grid patterns of the trapezoid rather than the regular grid patterns generated on the smallest rectangle enclosing the trapezoid t(58)=64.52, p<0.001, d=16.62, CI: 0.15; 0.16. **C.** The 50 eigenvector grid patterns used in this analysis.

**Supplemental Figure 3.**
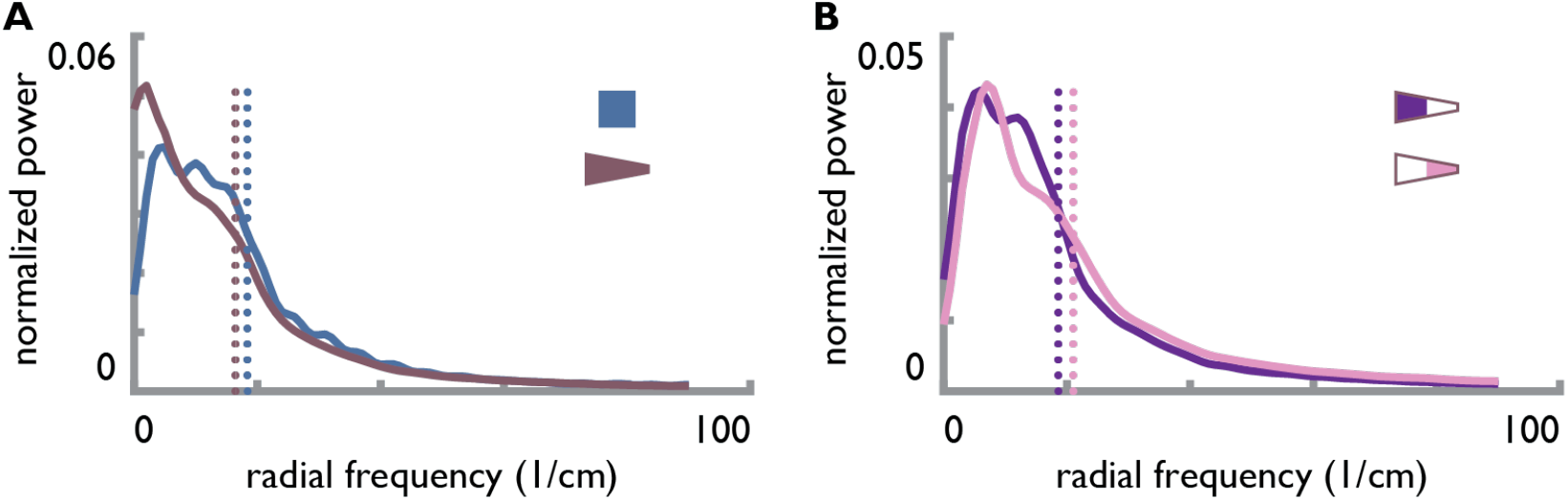
Spatial frequencies of eigenvector grid patterns. **A, B.** Radial power spectra based on two-dimensional FFT averaged across the 50 SR grid patterns. Average spatial frequencies were higher in the square than the trapezoid (**A**) and higher in the narrow compared to the broad part of the trapezoid (**B**). Dotted lines indicate mean radial frequencies.

**Supplemental Figure 4.**
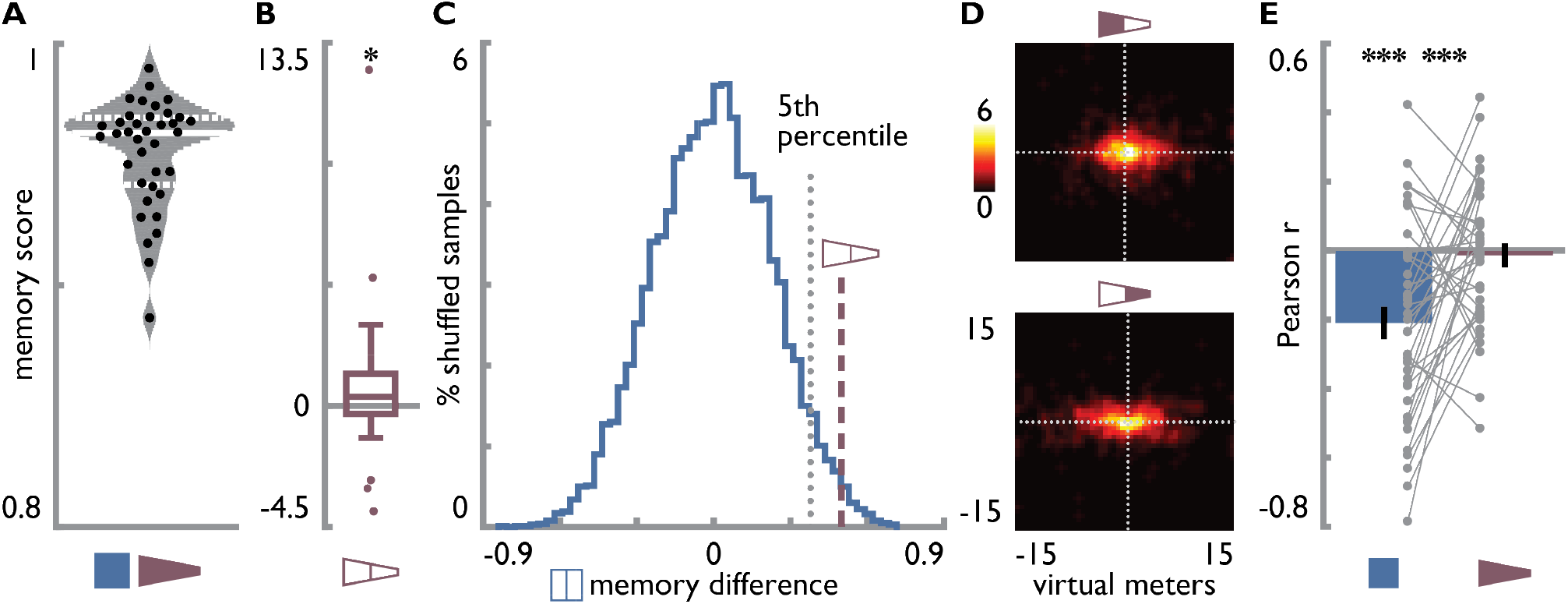
Positional memory. **A.** Distribution of average memory scores across participants. Grey area indicates normal kernel density estimate, solid white line shows median and dashed white lines show upper and lower quartile of distribution. Black circles show memory scores of individual participants. **B.** Positional memory error difference between the two parts of the trapezoid. Higher values indicate larger errors in the narrow part of the trapezoid. Data points more than 1.5 times the interquartile range above or below the upper or lower quartile were excluded as outliers (grey dots) for the main analysis, but comparable results are obtained without outlier exclusion (t(36)=1.50, p=0.020, d=0.25, CI: 0.04; 1.84). Boxplot represents median as well as upper and lower quartile of distribution, whiskers show most extreme value within 1.5 times the interquartile range from the upper and lower quartile respectively. **C.** The positional memory error difference observed between the trapezoid parts (dashed line represents mean difference across participants) was significantly lower than the critical value (5th percentile, dotted line) of a shuffle distribution (blue) obtained from computing error difference between the square halves across 10000 iterations. D. Heatmaps showing response locations for all trials across all participants for objects in the broad (top) and narrow (bottom) part of the trapezoid. Dotted lines show correct location in x- and y-dimension with their intersection representing the true position. **E.** Relationships between the distance to the closest boundary and the memory score were quantified using Pearson correlation. Correlation coefficients were consistently negative in the square, indicating better memory for positions closer to the wall. No such relationship was observed in the trapezoid and correlations differed between environments. * p<0.05 *** p<0.001

**Supplemental Figure 5.**
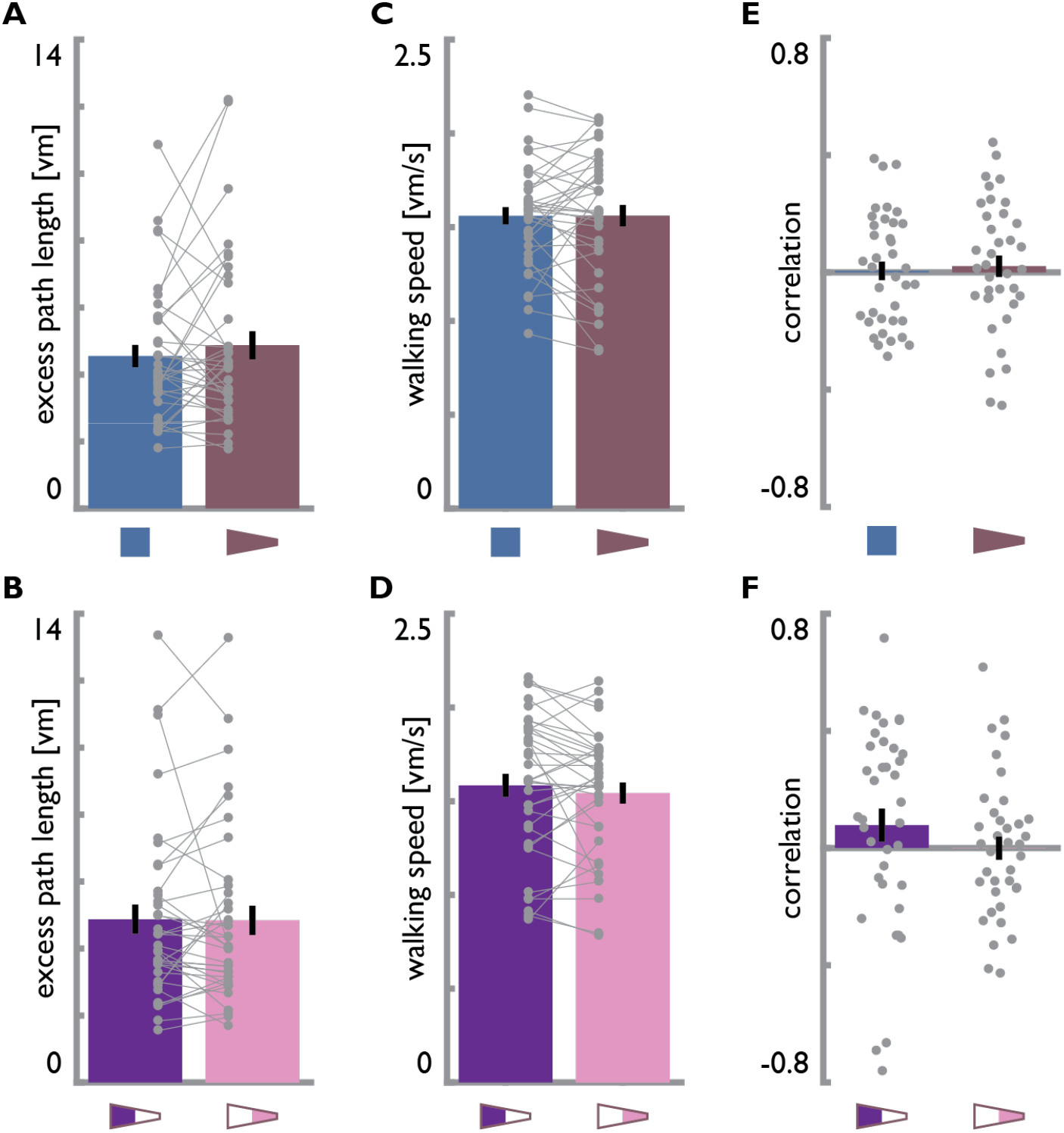
Navigation performance does not differ between environments. **A, B.** The excess path length of the trajectory from start to response position did not differ between (**A**) square and trapezoid or (**B**) the two parts of the trapezoid. **C, D.** Walking speed did not differ between (**C**) square and trapezoid or (**D**) the two parts of the trapezoid. **E, F.** Spearman correlation coefficients between the Euclidean distance from the start to the correct object positions and replacement error do not differ from 0 across subjects (**E**) in the square or trapezoid or (**F**) for objects located in the broad and narrow part of the trapezoid separately. Bars show mean±SEM and grey circles indicate individual subject data with lines connecting data points from the same participant.

**Supplemental Figure 6.**
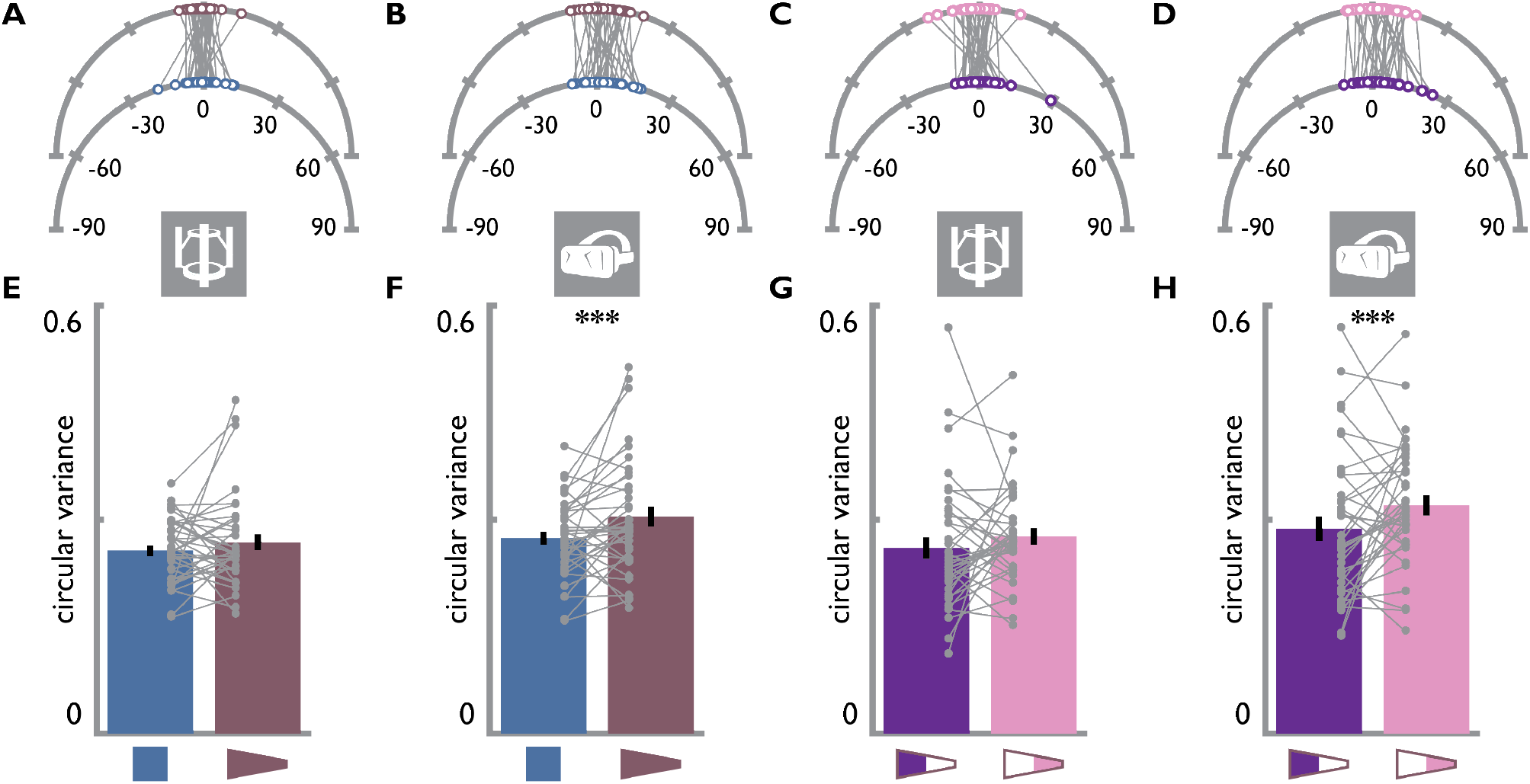
Head and body orientation during navigation. **A,B.** Circular means in degrees of (**A**) body and (**B**) head rotations centered on each trial’s direction from start to response position. Means are significantly clustered around 0° for both square and trapezoid and do not differ significantly. **C,D.** Circular means of (**C**) body and (**D**) head rotations centered on each trial’s direction from start to response position. Means are significantly clustered around 0 for trials with target object positions in the broad and narrow part of the trapezoid, respectively, and do not differ significantly. **E.** Circular variance of body rotations over trials averaged for each participant does not differ between square and trapezoid. **F.** Circular variance of head rotations over trials averaged for each participant is larger in the trapezoid than in the square. **G.** Circular variance of body rotations over trials averaged for each participant does not differ between navigation periods for target objects located in the broad or narrow portion of the trapezoid. **H.** Circular variance of head rotations over trials averaged for each participant is smaller when cued object position is in the broad compared to the narrow portion of the trapezoid. *** p<0.001

**Supplemental Figure 7.**
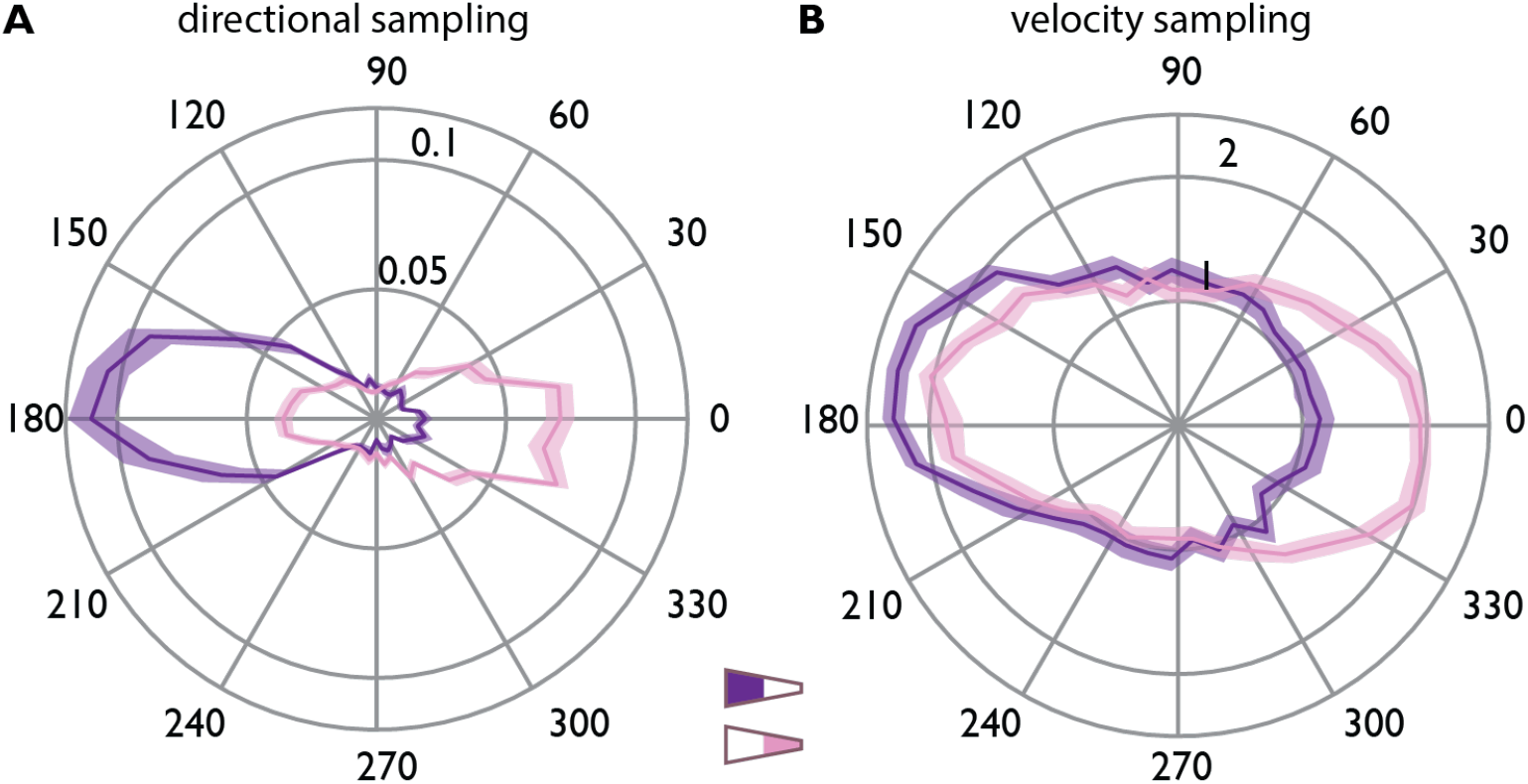
Angular and velocity sampling. **A.** Average angular sampling for 10° bins during navigation from a trial’s start position to the remembered object location. Radial axis shows proportion of time points facing in a directional bin. For trial’s targeting objects in the broad part of the trapezoid participants mostly faced towards the long base of the trapezoid (180°), whereas they more frequently faced towards the short base (0°) when targeting objects in the narrow part of the environment. **B.** Average movement speed (radial axis vm/s) for 10° directional bins for trials targeting objects in the broad and narrow part of the trapezoid. Navigation speed was higher along the long axis of the environment as indicated by higher movement speeds towards 0° and 180° for trials where participants targeted objects in the narrow and broad part of the trapezoid, respectively. In **A** and **B**, colored lines and shaded area show mean and SEM, respectively.

**Supplemental Figure 8.**
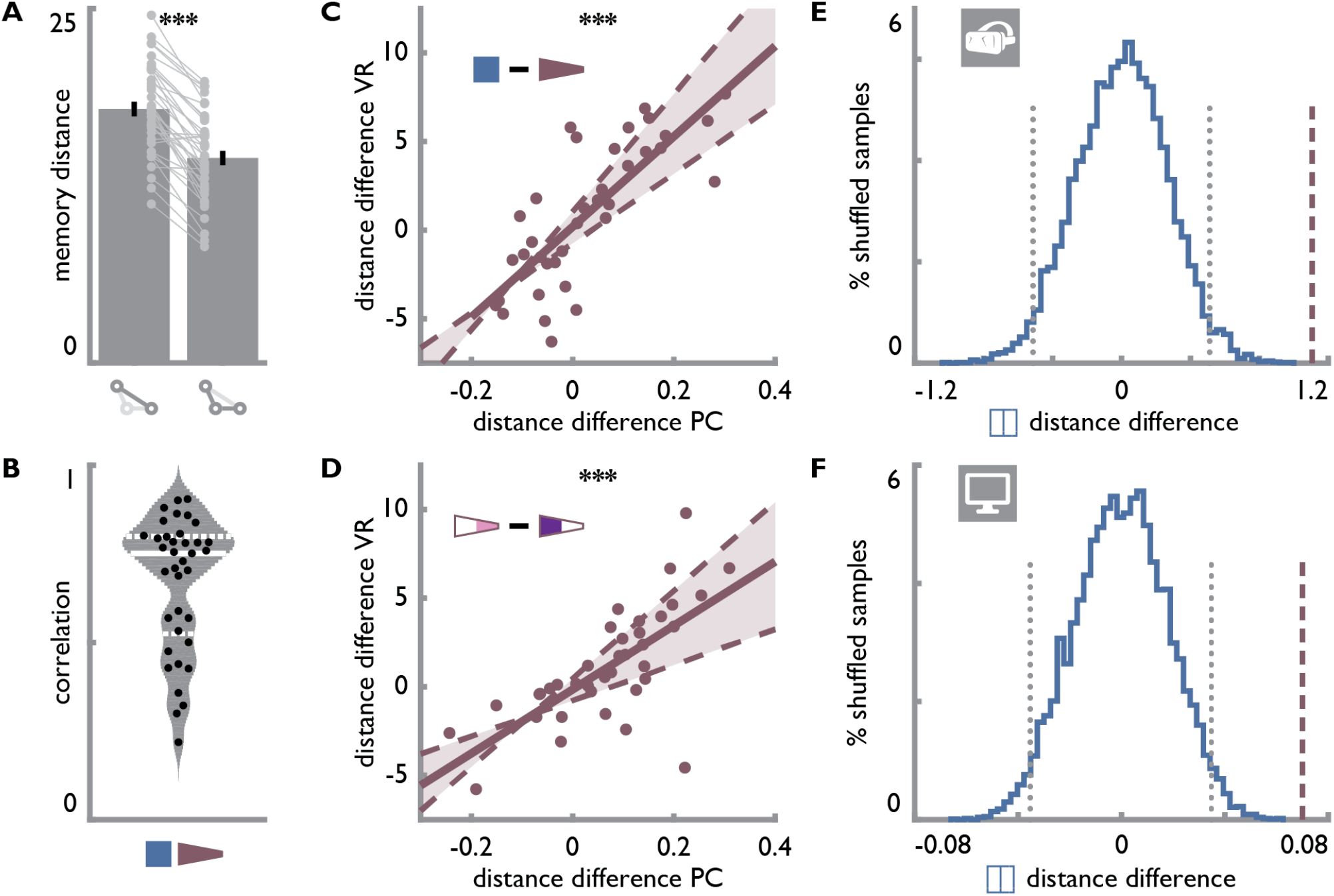
Distance estimates. **A.** Long distances (i.e. the base of the isosceles triangle formed by a triplet of positions) were estimated to be longer than the shorter distances (i.e. the legs of the isosceles triangle). Only within-triplet distances were estimated in VR. Bars show mean±SEM and grey circles indicate individual subject data with lines connecting data points from the same participant. **B.** Grey area indicates distribution of Spearman correlation (mean±SD r=0.69±0.19) coefficients between correct and estimated distances based on normal kernel density estimate. Solid white line shows median and dashed white lines show upper and lower quartile. Black circles show correlation coefficients of individual participants. **C.** The difference between distance estimates for identical distances in the square and the trapezoid was highly correlated between the computer screen and the VR version of the task. **D.** Significant correlation of distance difference between the two parts of the trapezoid obtained from distance estimates on the computer screen and in VR. Circles in **C** and **D** denote individual participant data; solid line shows least squares line; dashed lines and shaded region highlight bootstrapped confidence intervals. **E, F.** The distance difference observed between the trapezoid parts (dashed line) was more extreme than the critical values (dotted line) of the shuffle distribution (blue) obtained from computing the distance difference between the square halves across 10000 iterations for the distance estimates in VR (**E**) and on the PC (**F**).

**Supplemental Figure 9.**
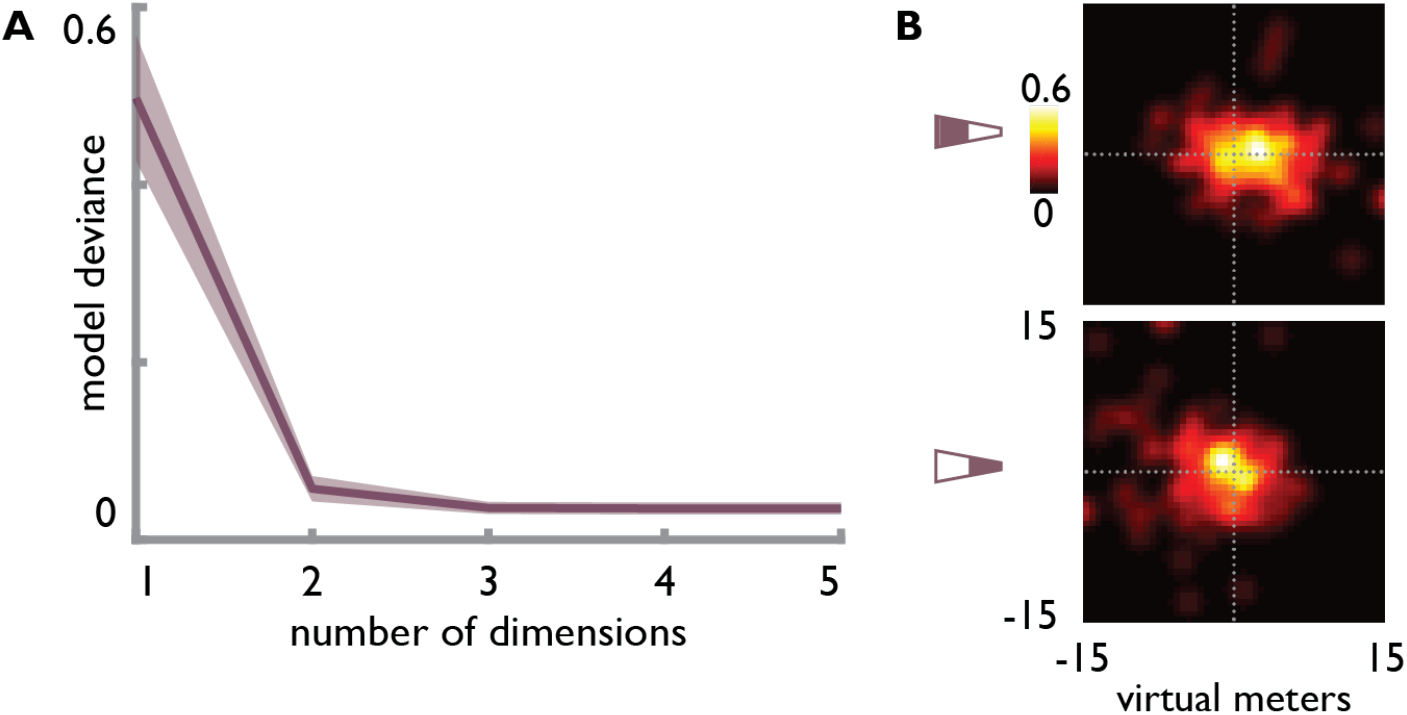
Two dimensions underlie distance estimates. **A.** Model deviance of GLMs using pairwise Euclidean distances of coordinates obtained from MDS to predict estimated distances for different numbers of dimensions (solid line shows mean model deviance across participants, shaded area indicates SEM). In line with our a priori assumption that two dimensions underlie the distance estimates, model deviance sharply drops when using two rather than one dimension and there is no substantial benefit from including three or more dimensions. **B.** Heatmaps showing positions reconstructed using multi-dimensional scaling and Procrustes transform for objects in the broad (top) and narrow (bottom) part of the trapezoid. Dotted lines show correct position in x- and y-dimension with their intersection representing the true position.

**Supplemental Figure 10.**
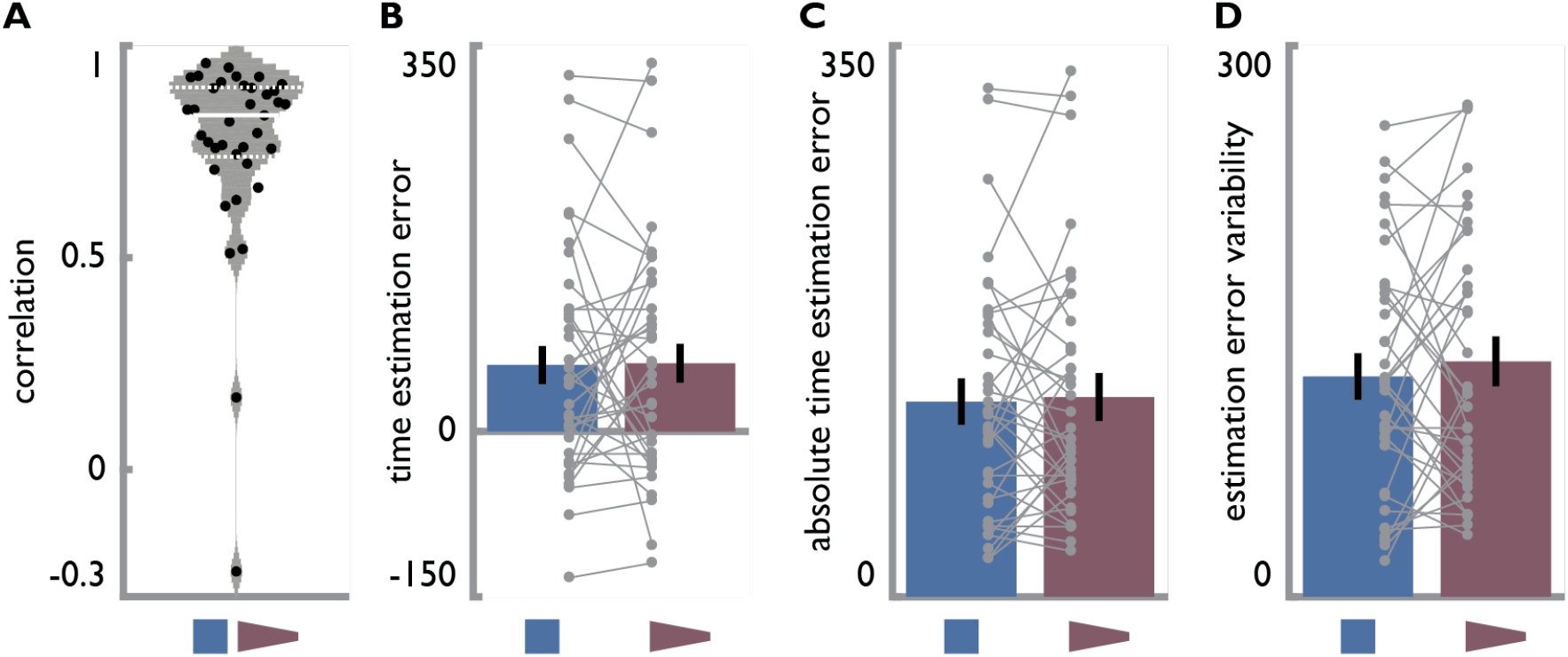
Time estimates do not differ between environments. **A.** Grey area indicates distribution of Spearman correlation coefficients (mean±SD r=0.77±0.23) between true and estimated times based on normal kernel density estimate. Solid white line shows median and dashed white lines show upper and lower quartile. Black circles show correlation coefficients of individual participants. **B.** Averaged time estimation errors did not differ between the two environments. C. Averaged absolute time estimation errors did not differ between the two environments. D. The variability of time estimates as measured by their standard deviation did not differ between environments. Bars show mean±SEM and grey circles indicate individual subject data with lines connecting data points from the same participant.

## Acknowledgments

We thank Jackeline Neves Pereira for pilot work leading up to the final experimental design. CFD’s research is supported by the Max Planck Society; the European Research Council (ERC-CoG GEOCOG 724836); the Kavli Foundation, the Centre of Excellence scheme of the Research Council of Norway – Centre for Neural Computation, The Egil and Pauline Braathen and Fred Kavli Centre for Cortical Microcircuits, the National Infrastructure scheme of the Research Council of Norway – NORBRAIN; and the Netherlands Organisation for Scientific Research (NWO-Vidi 452-12-009; NWO-Gravitation 024-001-006; NWO-MaGW 406-14-114; NWO-MaGW 406-15-291). The funders had no role in study design, data collection and analysis, decision to publish or preparation of the manuscript.

## Author contributions

J.B., C.B. and C.D. conceived the experiment. J.B., T.R., M.N. and C.D. designed the experiment. T.R. collected the data. J.B. analyzed the data and wrote the manuscript, with input from M.N., C.B. and C.D. W.C. performed the model analysis under supervision of C.B. All authors discussed the results and contributed to the final manuscript.

